# Exponential trajectories, cell size fluctuations and the adder property in bacteria follow from simple chemical dynamics and division control

**DOI:** 10.1101/487504

**Authors:** Parth Pratim Pandey, Harshant Singh, Sanjay Jain

## Abstract

Experiments on steady state bacterial cultures have uncovered several quantitative regularities at the system level. These include, first, the exponential growth of cell size with time and the balanced growth of intracellular chemicals between cell birth and division, which are puzzling given the nonlinear and decentralized chemical dynamics in the cell. We model a cell as a set of chemical populations undergoing nonlinear mass action kinetics in a container whose volume is a linear function of the chemical populations. This turns out to be a special class of dynamical system that generically has attractors in which all populations grow exponentially with time at the same rate. This explains exponential balanced growth of bacterial cells without invoking any regulatory mechanisms and suggests that this could be a robust property of protocells as well. Second, we consider the hypothesis that cells commit themselves to division when a certain internal chemical population reaches a threshold of N molecules. We show that this hypothesis leads to a simple explanation of some of the variability observed across cells in a bacterial culture. In particular it reproduces the adder property of cell size fluctuations observed recently in *E. coli*, the observed correlations between interdivision time, birth volume and added volume in a generation, and the observed scale of the fluctuations (CV ~ 10-30%) when N lies between 10 and 100. Third, upon including a suitable regulatory mechanism that optimizes the growth rate of the cell, the model reproduces the observed bacterial growth laws including the dependence of the growth rate and ribosomal protein fraction on the medium. Thus, the models provide a framework for unifying diverse aspects of bacterial growth physiology under one roof. They also suggest new questions for experimental and theoretical enquiry.

## I. INTRODUCTION

The simplest cells, bacteria, exhibit several generic phenomena that were discovered decades ago but still require explanation. One such phenomenon is that in steady state bacterial cultures the size of an individual bacterial cell grows exponentially with time between birth and division. This was observed early on in [1, 2] and its existence in various organisms has been the subject of debate [3, 4]. Recently detailed single cell experiments [5–11] have confirmed this property for many bacterial species in steady state cultures. Intracellular molecular populations have also been observed to grow exponentially in *Escherichia coli* cells within a generation [12]. Since bacterial cells divide, the range of size over which exponential trajectories are seen is limited to a factor of two (or similar), thereby potentially permitting alternate fits to the data. However, the exponential function fits quite well and is therefore at least a very good approximation. In other single-celled organisms such as *Schyzosaccharomyces pombe* a clear departure from exponential trajectories is seen [4, 13], so this is not a property that can be taken for granted. Why most bacteria exhibit this property remains an unanswered question.

In linear autocatalytic systems [14–17] exponential growth is not surprising. The asymptotic rate of exponential growth of chemical populations in such systems is the largest eigenvalue of the matrix defining the dynamical system. However, the chemical dynamics of the intracellular molecular species in a bacterial cell is highly nonlinear and therefore exponential trajectories are surprising. In this paper we provide an explanation for how cells can exhibit exponential trajectories of intracellular chemicals and cell volume in spite of the nonlinearity of the dynamics. We show that ordinary differential equations representing mass-action based nonlinear chemical dynamics in an expanding container whose volume is linearly dependent on the constituent populations, have a special scaling property or ‘quasi-linearity’. Such nonlinear systems naturally have exponentially growing trajectories as attractors.

Another unexplained phenomenon is ‘balanced growth’ [18], the remarkable coordination between thousands of intracellular chemicals so that all of them (on average) double in the same time. This is required for self-replication: a cell at birth must grow in such a way that division, when it happens, produces two daughters identical to itself; hence the mother-at-division must have twice of everything as the daughter-at-birth. In spite of their decentralized dynamics characterized by reaction-specific catalysts and rate constants, how do thousands of chemicals conspire to double at the same time? It is normally supposed that regulatory mechanisms involving checkpoints and feedback loops are responsible for this coordination. We argue that this sophistication is not needed. The exponential trajectories that we find as generic attractors in autocatalytic systems have all chemical populations increasing at the same exponential rate even when no regulation is present. Thus their ratios are automatically constant in time once they reach the attractor. The genericity and robustness of these attractors suggests that balanced growth could have been a property of protocells at the origin of life. The protocell literature has been concerned with this question [19–24] and has been an inspiration for this work, though the solution presented in this paper is different from the one advocated there. At a mathematical level we show that the growth rate as well as the ratios of chemicals in the attractor are determined from a nonlinear generalization of the eigenvalue equation of a matrix.

Another phenomenon that has consistently received attention but is still not understood is the origin and scale of variation in interdivision time and cell size; reviewed in [25–28]. Genetically identical bacterial cells subjected to the same steady environment exhibit a phenotypic variation in both these quantities. In the steady state the volume of the cells at birth and the time between birth and division have steady distributions with a constant mean and standard deviation (s.d.) and a coefficient of variation, CV = s.d./mean, in the range of 10-30%. The mean size of the cell-at-birth can be changed by close to a factor of 10 by choosing different environmental conditions; its CV remains between 10-30% [9, 11, 29]. The CV of the molecular population of an abundant intracellular chemical across cells in a bacterial culture (the extrinsic CV) is also 10-30% [30–32]. It is believed that the source of variation lies in the process that controls cell division; however the molecular basis of this process is not fully understood, even for *E. coli*, and nor is the origin of the scale of CV. In this paper we explore, through a mechanistic mathematical model, the hypothesis [33] that division is controlled by the intracellular population (not concentration) of a certain molecule reaching a fixed threshold ~ *N*, a parameter of the model. Then N controls both the average size and the fluctuations in size and interdivision time. We provide an explicit expression for the average cell volume *V* in terms of *N* and the other molecular parameters of the model. Fluctuations arise because molecular production is stochastic and hence the interdivision time, which under this hypothesis is the first passage time for this molecule to reach its threshold, is stochastic. The CV of the interdivision time is 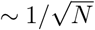; hence the CV of cell volume and of extrinsic molecular populations is also 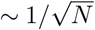. We note that this hypothesis therefore explains the observed range 0.1-0.3 of the CV of all these quantities provided *N* lies between 10 and 100. We compare the theoretical distributions of size, interdivision time and added volume with the experimental data of [11] to constrain the range of *N* for *E. coli*.

Size fluctuations in *E. coli* have been recently observed [10, 11] to satisfy the ‘adder’ property [34, 35], wherein the volume added by a cell in each generation is independent of its birth volume in that generation. This property has been explored in various phenomenological models [8, 10, 11, 34–36] and implies specific correlations between the cell volume at birth and division, and between interdivision time and birth volume. It has also been shown to arise in a linear mechanistic model [37]. We show that the adder property appears robustly for a wide class of non-linear mechanistic models under the hypothesis that division is triggered when a molecular population reaches a threshold value, generalizing the results of [37]. We indicate circumstances where fluctuations depart from the adder.

Fluctuations of the growth rate across cells in a steady state culture have been measured in the literature and CV values ranging from 10-40% have been reported [11, 12, 38, 39]. Our models suggest that the physical origin of fluctuations in the growth rate is different from that of interdivision time and cell size. The simplest and most natural models that we consider exhibit a smaller CV (1-5%) than that reported in the experimental literature, though larger values can also be accounted for. Our models also reproduce the observed cross-over in the CV of an intracellular molecular population as a function of the mean population 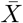 (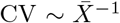 for small 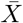 and constant for large 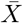) [31].

Growth laws of bacterial composition [40–43] constitute another class of generic phenomena in bacteria which describe how the cellular steady state growth rate and ribosomal protein fraction in the cell depend upon the medium. These bacterial growth laws have received a substantial theoretical attention [43–50] and are much better understood compared to the phenomena mentioned in the previous paragraphs. A particular model that we discuss in detail reproduces the bacterial growth laws in addition to the other phenomena mentioned above. Thus this work provides possible explanations and a unified understanding of a number of generic properties of bacteria, including exponential trajectories in the steady state, balanced growth of cells, the growth laws of bacterial composition, as well as the scale and correlations of the fluctuations of several cellular variables. The work makes predictions that can be experimentally tested.

### Organization of the paper

In section II we describe the general class of mathematical models of cells that we consider. In section III a specific nonlinear model is considered and results based on numerical simulations are presented that reproduce the features mentioned above. In addition to intrinsic stochasticity in chemical dynamics we also discuss here the effect of stochasticity in the partitioning of chemicals at division and stochasticity in the threshold value of the division trigger. In section IV we derive analytically several of the results presented in section III. In particular we provide an analytic understanding of the robustness (or universality) of the distributions of cell size, intracellular populations and interdivision time, and the non-universality of the distribution of the cellular growth rate. Section V discusses the mathematical and physical basis for exponential trajectories and balanced growth and shows that these are generic properties of mass action chemical dynamics in expanding containers whose volume is a linear function of the internal chemical populations. This section also extends the analytical results of section IV to a large class of models. Section VI discusses the unification of the bacterial growth laws with the other cellular properties. Finally section VII gives a detailed summary of the model assumptions and results and discusses future experimental and theoretical directions. The supplementary material contains additional figures and proofs. Some readers may benefit by turning directly to the summary in section VII for a quick guide to the main results including equations and figures.

## II. A GENERAL MODEL OF A GROWING-DIVIDING CELL

We model the cell as a container with *n* + 1 chemical species, whose molecular populations at time *t* are denoted *X_i_*(*t*), *i* = 1, 2,…, *n*, and *Z(t)*. *Z* is the species triggering cell division and **X** = (*X*_1_,…, *X_n_*) represents all the other chemicals in the cell. The population dynamics is given by

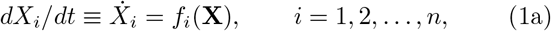

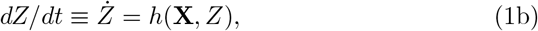

where the functions *f_i_* and *h* encapsulate the kinetics of the chemical reactions in the cell. The *Z* population is assumed to have a negligible effect on the **X** dynamics; hence the *f_i_* are independent of *Z*. The volume of the cell is assumed to depend linearly on the 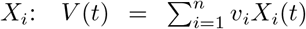 where the *v_i_* are constant parameters of the model. Again, the contribution of *Z* to *V* is assumed to be negligible. Explicit examples of the functions *f_i_* and h will be discussed later.

### Division control

Starting from some initial condition for the *n* + 1 populations, the populations are evolved according to (1). The function *h* is assumed to be positive, hence *Z* increases under (1). When *Z* reaches the threshold *Z_c_*, the cell divides. The two daughter cells are assumed to be identical, hence each gets half of every chemical in the cell. Since in the model we track one of the two daughters, at division all *n* + 1 populations are replaced by half their values. This defines the state of the daughter at birth in the next generation and with that initial condition for the populations the above procedure is iterated.

### Resetting of *Z* after triggering; delay between triggering and division

The above defines the sim plest version of the model. It is useful, however, to consider two generalizations motivated by empirical observations. The first involves a resetting of the *Z* population after it triggers division. In the above scheme, *Z_b_*, the value of *Z* in a daughter at birth is always *Z_c_*/2. However specific biochemical processes may cause a degradation of the triggering molecule on a short time scale after triggering of division. In order to model this we assume for simplicity that when *Z* reaches *Z_c_* it triggers division and is also instantaneously reset to a value *Z_r_* ≤ *Z_c_*. *Z_r_* is another parameter of the model; *Z_r_* = 0 implies complete degradation of the triggering molecule; *Z_r_* = *Z_c_* implies no degradation. Several properties that we discuss (e.g., the adder property) hold for the entire range of values 0 ≤ *Z_r_* ≤ *Z_c_*. The second generalization is that division follows triggering (and resetting of *Z*, if any) after a time delay *τ*_1_, which is another parameter of the model. When *τ*_1_ = 0, then division immediately follows triggering and resetting; hence *Z_b_* = *Z_r_*/2. When *τ*_1_ > 0 we assume that after triggering (and instantaneous resetting of *Z* to *Z_r_*) all *n* + 1 populations continue to evolve via (1) for a fixed time *τ*_1_, whereupon cell division takes place and all are halved. These generalizations are motivated by evidence that the replication of DNA is initiated when a certain protein reaches a threshold population, and this chemical degrades soon after the initiation of replication to avoid multiple replication rounds. Cell division follows the initiation of DNA replication after the lapse of a certain time.

To summarize, the population dynamics in the model is a combination of continuous dynamics described by (1) and discrete events. The **X** variables evolve via (1a) from their initial values upto the time of cell division; only at division do they experience a discrete change: they are halved. The *Z* variable evolves via (1b) from its initial value upto the point where it reaches the threshold *Z_c_*. There it is instantaneously reset to *Z_r_*. Subsequently it again evolves via (1b) for a fixed time *τ*_1_. That defines the time of cell division. At that time *Z* is halved (along with the *X_i_*). That brings the cell to the beginning of the next generation and a new set of initial values of the *n* +1 populations. Thereupon the procedure is iterated.

Note that the dynamics of *Z* explicitly depends upon the *X* sector through *h*(**X**, *Z*). On the other hand, *Z* does not affect the short time-scale dynamics of the *X* sector since *f_i_* are assumed to be independent of *Z*. However *Z* exercises a discrete control over the *X* variables through the triggering of division, which in effect determines how large the *X_i_* can grow before cell division occurs and they halve. The functions *f_i_* contain all the interactions among the *X* chemicals and encapsulate most of the complexity of the intracellular dynamics.

The above formulation describes the deterministic version of the the model. We also consider various sources of stochasticity. One is the intrinsic stochasticity in the chemical dynamics resulting from the populations being non-negative integers instead of continuous variables and each reaction having a certain probability of occurring. Then Eqn. (1) is replaced by its stochastic counterpart. Further at division the chemicals may not partition equally into both daughters. Also *Z_c_*, the threshold value of of *Z* at which division is triggered may vary from generation to generation. The specific implementation of these sources of stochasticity in the model will be discussed later.

## III. A SPECIFIC EXAMPLE WITH 3+1 CHEMICAL SPECIES

We now consider a concrete example in which the *X* sector is described by the Precursor-Transporter-Ribosome (PTR) model discussed in [50] containing *n* = 3 species. Later on we will argue that several of the results of the PTR model hold for the general model described above with a very broad class of functions *f_i_* and *h*. Here *X*_1_ = *P* is a coarse-grained population variable representing the total number of amino acid molecules (precursors or monomers out of which proteins are constructed), *X*_2_ = *T* is the total number of transporter and other metabolic enzyme molecules in the cell responsible for making *P* from the food molecules available in the extracellular medium, and *X*_3_ = *R* is the total number of ribosomes in the cell (catalysts for making *T* and *R* from the *P* molecules). The equations of the *X* sector (or PTR sector) are

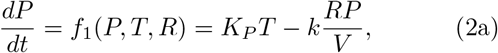

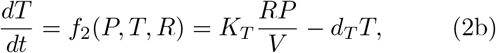

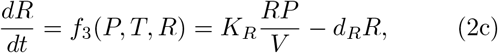

where

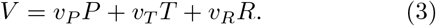

The constants *K_P_, K_T_, K_R_, k, d_T_, d_R_, v_P_,v_T_, v_R_* are parameters of the model. The rate constant *K_P_* represents the efficiency of metabolic enzymes *T* in transporting and producing *P* from external food. It is an increasing function of the external food concentration [*F*] and the quality of the food source *q* (the number of *P* molecules produced per food molecule transported in). *k* represents ribosomal catalytic efficiency, and is the peptide elongation rate per unit concentration of *P*. *d_T_*, *d_R_* are degradation rates. *v_P_, v_T_, v_R_* define the contributions of the individual species to the cellular volume *V*. It is convenient to parametrize the rate constants *K_T_*, *K_R_* as follows:

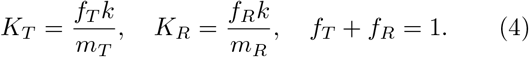

Here *f_T_* and *f_R_* represent the fractions of ribosomes engaged in making *T* and *R* respectively, and *m_T_* and *m_R_* are the number of amino acid residues (or units of *P*) in a *T* molecule and ribosome respectively. It follows that the mass of the PTR cell (in units of mass of a *P* molecule) is *M* = *P* + *m_T_T* + *m_R_R*. For more details see [50].

In the *Z* sector the function *h* is defined by

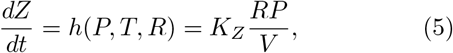

where *K_Z_* is a constant. This form assumes that the production of the molecule that triggers cell division mirrors the growth of the cell (this term has the same form as the growth term of *T* and *R* in (2)). Note that in (5) *h*(**X**, *Z*) is independent of *Z*. We will later (in section IV D) give results when h has a non-trivial *Z* dependence arising from auto-regulation of *Z* (e.g., cooperativity). The PTR model [50] has been independently extended by [51] to include a sector whose dynamics is given by (5) and that triggers division upon reaching a population threshold. Their treatment differs from ours in various respects, in particular, that the new sector affects the local dynamics of the PTR sector through the volume. They have also observed the adder property of the cell volume in numerical simulations. Apart from that commonality, ref. [51] and the present work explore different aspects of the models.

### A. Deterministic version; numerical results

The rules mentioned above fully specify the deterministic dynamical system once the parameters of the PTR sector (*K_P_, k, d_T_, d_R_, m_T_, m_R_, f_R_, v_P_, v_T_, v_R_*), the *Z* sector (*K_Z_, Z_c_, Z_r_*) and *τ*_1_ are specified. For simplicity we consider the case where *τ*_1_ = 0 (*τ*_1_ > 0 is discussed later). Thus, when Z reaches Z_c_, the cell divides without any delay, i.e., the populations *P, T, R* are halved, and the birth value of *Z* is simultaneously set to *Z_r_*/2. Subsequently the populations follow Eqs. (2) and (5) until *Z* again reaches *Z_c_* and the procedure is repeated. Simulations are shown in Fig. 1. Most parameters of the PTR sector are chosen in the ballpark of realistic values [42, 43, 52, 53] and reproduce known experimental data on *E. coli* within a factor of order unity (namely, the values of *P, T, R* and the growth rate). The values of *v_T_* and *v_R_* have been set equal to *v_P_* for simplicity; we are not aware of experiments that measure the *v_i_* independently. The value of *f_R_* has been chosen so as to maximize the steady state growth rate of the PTR cell with the above mentioned choice of parameters; this procedure reproduces the bacterial growth laws as discussed in [50]. In the *Z* sector, the value of *Z_c_* and *Z_r_* are chosen to get the scale of size fluctuations in the experimentally observed ballpark (it will be seen that only the combination *Z_c_* − *Z_r_*/2 matters). *K_Z_*, an unknown parameter of the model is chosen so that the steady-state birth volume of the cell turns out in the correct ballpark ~ 1(*μm*)^3^. Our conclusions are robust to independent variation of each parameter (within collective limits as discussed later).

**FIG. 1:**
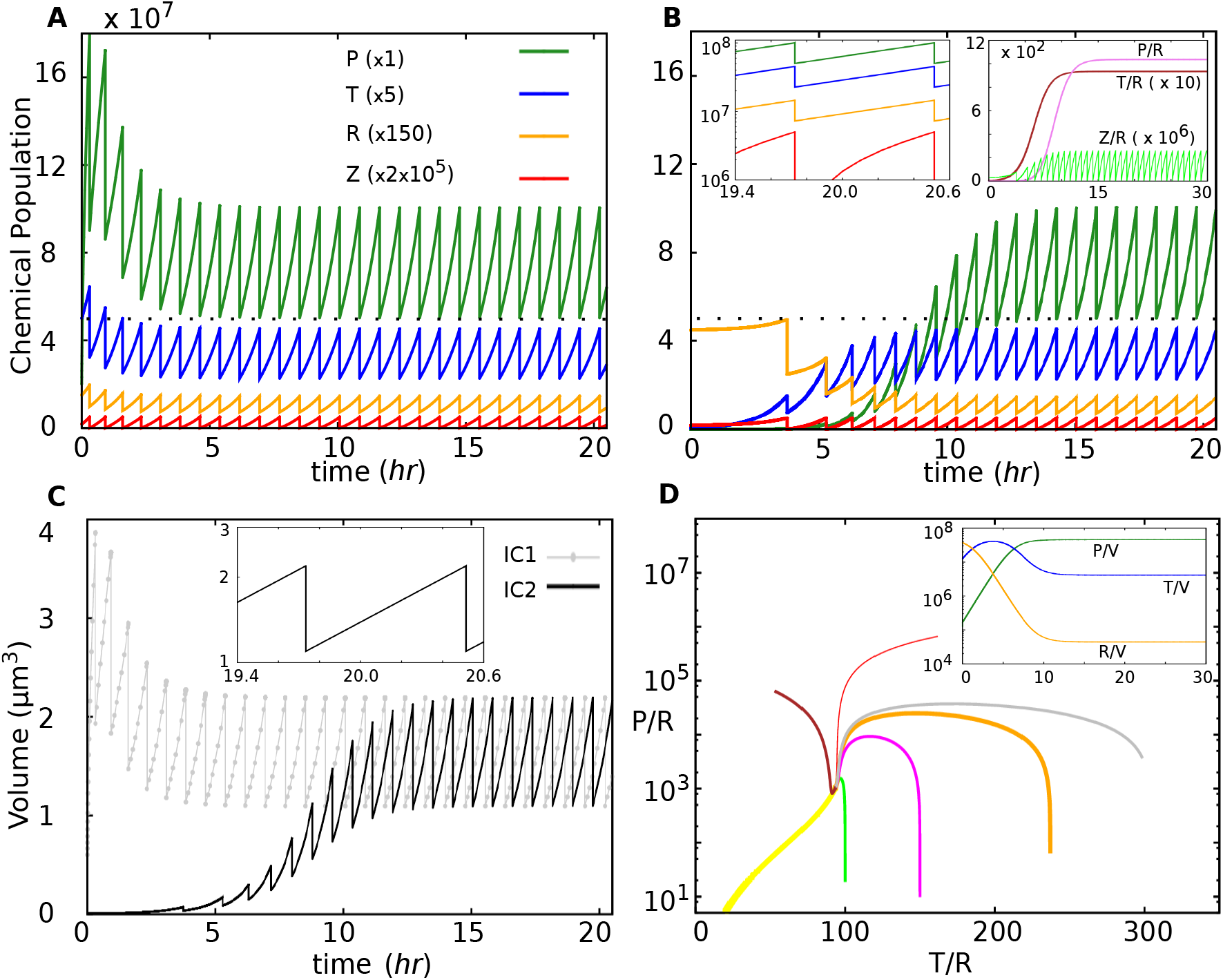
Plot of the chemical populations *P, T, R, Z* and volume *V* vs time *t* for deterministic trajectories. Parameters of the model are *K_P_* = 500 *hr*^−1^, *k* = 10^−3^ *hr*^−1^ (*μm*)^3^, *d_T_* = 0.1 *hr*^−1^, *d_R_* = 0, *m_T_* = 400, *m_R_* = 10000, *v_P_* = *v_T_* = *v_R_* = 2 * 10^−8^(*μm*)^3^, *f_R_* = 0.19377, *K_Z_* = 10^−11^ *hr*^−1^ (*μm*)^3^, *Z_c_* = 25, *Z_r_* = 0, *τ*_1_ = 0. **A** The trajectory of a cell with the initial conditions IC1: *P*(0) = 2 * 10^7^, *T*(0) = 10^7^, *R*(0) = 10^5^, *Z*(0) = 9, and **B** IC2:*P*(0) = 10^3^, *T*(0) = 10^5^, *R*(0) = 3 * 10^5^, *Z*(0) = 9. The population values in **A** and **B** have been multiplied by the factors mentioned in the legend of **A** to bring them on the same figure. The dynamics converges to a steady state (a periodic solution or limit cycle attractor). The chemical populations in the daughter cell at birth in the steady state are *P_b_* = 5.023 * 10^7^, *T_b_* = 4.529 * 10^6^, *R_b_* = 4.844 * 10^4^ (*P_b_* is indicated by a black dotted line in **A** and **B**). The period, or inter-division time *τ* = 0.781 *hr*. (C) The volume of the cell, *V*, defined by Eq. (3), as a function of time for the two initial conditions IC1 and IC2. In this and subsequent figures, the unit of the time axis is hr and the volume axis (*μm*)^3^. First inset of **B** and inset of **C**: Semilog plots of *P, T, R, Z* and *V* vs time in the steady-state with IC2. The plots of *P, T, R* and *V* are piecewise linear and have the same slope 0.888 *hr*^−1^ (slope of the natural logarithm of the quantity vs time), showing that they all increase exponentially with the same rate in the growth phase of the limit cycle attractor. The slope agrees with formula (8c). Second inset of **B** plots the ratios of chemical populations as a function of *t* on the trajectory starting from IC2. The ratios of populations of the PTR sector become constant on the limit cycle attractor and the constant values agree with Eqns. (8a),(8b). **D** Simulated phase portrait of trajectories from diverse initial conditions projected on to the 2-dimensional space of *T*/*R* and *P*/*R* showing that the trajectories converge to the same attractor independent of initial conditions. Inset of **D**: Time dependence of concentrations of *P, T, R* on the trajectory starting from IC2, becoming constant on the limit cycle attractor.

As evident from Fig. 1, after a transient that depends on initial conditions, the dynamics converges to a steady state. This steady state is a limit cycle attractor in which the just born daughter cell grows to twice its size (the populations *P, T, R* and the cell volume double) and then divides into two halves bringing the system back to the same daughter state. It is seen that in the attractor, between birth and the next division (i.e., within a single generation of the cell) the chemicals of the X sector grow exponentially with time at the same rate (inset of Fig. 1B). The same is true of *V* (inset of Fig. 1C), a consequence of (3). Because of exponential growth the ratios *T*/*R* and *P*/*R* remain constant in time throughout the attractor. Hence the concentrations of the X sector chemicals also remain constant in the attractor (inset of Fig. 1D). During the transient period before the attractor is reached the ratios are not constant. For all the initial conditions considered (with 0 ≤ *Z*(0) < *Z_c_*) the trajectory converges to the same steady state (Fig. 1D), suggesting that the limit cycle is a stable attractor with a wide basin of attraction. When the parameters are varied within a wide range (specified later; section IV A) the same dynamical behaviour is observed: initial transient leading to a limit cycle attractor characterized by exponential trajectories in the growth phase and constant ratios of chemicals.

### B. PTRZ with stochastic chemical dynamics; numerical results

In Fig. 2A we show a typical trajectory of the PTRZ cell following stochastic chemical dynamics. The difference from the deterministic version is that the evolution of *P, T, R, Z* is no longer given by the differential equations (2) and (5) but by a stochastic version of (2) and (5). The populations are now non-negative integers and are updated according to the probabilities of the chemical reactions, the latter being proportional to the corresponding terms in the differential equations. The rest of the dynamical rules are the same as before (starting from any given initial condition, the populations *P, T, R, Z* are evolved with time until *Z* = *Z_c_*, at which point *P, T, R* are halved (and rounded off to the nearest integer value), *Z* is set to *Z_r_*/2 and the procedure iterated). No new physical parameter is introduced in the stochastic dynamics. The algorithm used for numerical simulations is mentioned in Computational Methods below.

**FIG 2.**
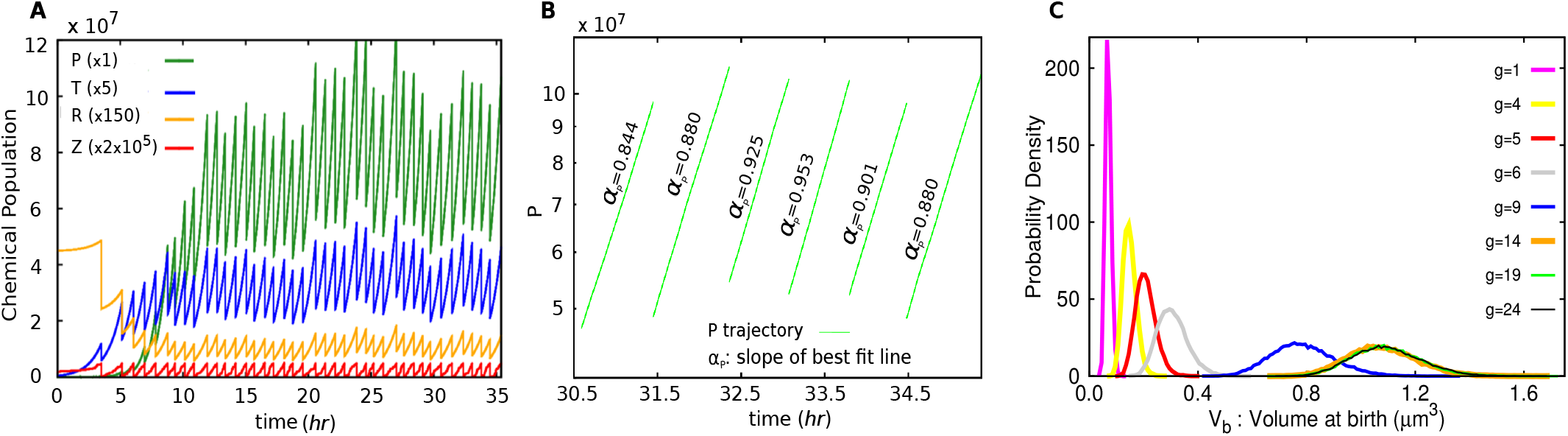
Effect of stochasticity in chemical dynamics of the PTRZ cell. Parameter values are as in Fig. 1. **A** A stochastic trajectory of *P, T, R.Z* starting from IC_2_ (see legend of Fig. 1). **B** *P* vs time *t* in a semi-log plot for five consecutive generations in the statistical steady state. The slope *α^P^* of the best fit straight line (ln *P* vs *t*) varies from generation to generation. **C** Time evolution of the volume distribution of 30000 cells at birth. The distribution at the beginning of the *g*^th^ generation (or just after *g* divisions) is shown for various values of *g*. In this case the initial distribution of *V* is a *δ*-function at *V* = 0.06 (*μm*)^3^, since all the trajectories start at *t* = 0 from the initial condition IC3: *P*(0) = *T*(0) = *R*(0) = 10^6^, *Z* = 0. The distributions at *g* = 19 and 24 are indistinguishable, suggesting a stationary state.

#### 1. Stochastic steady state

We observe in Fig. 2A that after a few rounds of growth and division the trajectory of the cell reaches a statistical steady state, a stochastic version of the deterministic steady state seen in Fig. 1A,B). In the stochastic steady state the daughter cells produced at the beginning of each cycle are no longer identical from generation to generation (in terms of the values of *P, T, R, V_b_*) as was the case in the deterministic simulation, and *τ*, the interdivision time also varies from cycle to cycle. The latter variation arises because the first passage time of *Z* to reach *Z_c_* varies from cycle to cycle. Fig. 2B shows that in the statistical steady state the population growth of the *X* sector chemicals within a generation can be approximated by an exponentially growing trajectory with the effective growth rate varying from generation to generation.

We performed 30000 stochastic simulations for the PTRZ model each with the same initial condition as described in the caption of Fig. 2C. The cellular variables were tracked for each trajectory. The distribution of the cell volume at birth, *V_b_*, across the 30000 trajectories at different generations is shown in Fig. 2C. The distribution becomes stationary after a few generations. The same steady state distribution is obtained by (a) starting from a different initial distribution of the 30000 cells and (b) sampling over different generations of a single trajectory after the initial transient period. The distributions of *V_d_, P_b_, T_b_, R_b_, τ* and *α* are also found to converge to their respective steady state distributions independent of the initial distribution of cellular configurations (*α* is the ‘growth rate’ of the cell in a generation, defined as the slope of the best straight line fit of ln *V*(*t*) vs *t* in that generation). This indicates that the statistical steady state is a stable attractor of the dynamics characterized by fixed distributions of the cellular variables.

#### 2. Distributions of cell size, interdivision time and growth rate, correlations and the adder property

The steady state distributions are shown in Fig. 3. The averages of the steady state distributions are: 〈*P*〉 = 5.012 * 10^7^, 〈*T*〉 = 4.518 * 10^6^, 〈*R*〉 = 4.833 * 10^4^, 〈*V_b_*〉 = 1.094*μm*^3^, 〈*V_d_*〉 = 2.190*μm*^3^, 〈Δ〉 = 1.096*μm*^3^, 〈*τ*〉 = 0.782 *hr*, 〈*α*〉 = 0.888 *hr*^−1^. These are close to the deterministic values in Fig. 1. The model predictions of the distributions of the above quantities rescaled by their means (except the rescaled *α* distribution) are largely independent of all the PTRZ model parameters (and hence independent of growth rate, ratios of chemicals, etc.) and of *K_Z_* when *τ*_1_ = 0. They only depend upon the quantity *N* = *Z_c_* − (1/2)*Z_r_*. (See Fig. S1 for a completely different parameter set giving the same rescaled distributions.) The reason for this extraordinary robustness will be discussed later. We find that the CV of the Δ, *V_b_, V_d_* and *τ* distributions are consistent with

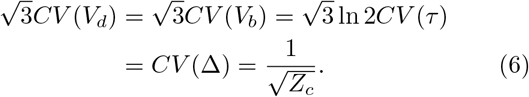

**FIG 3.**
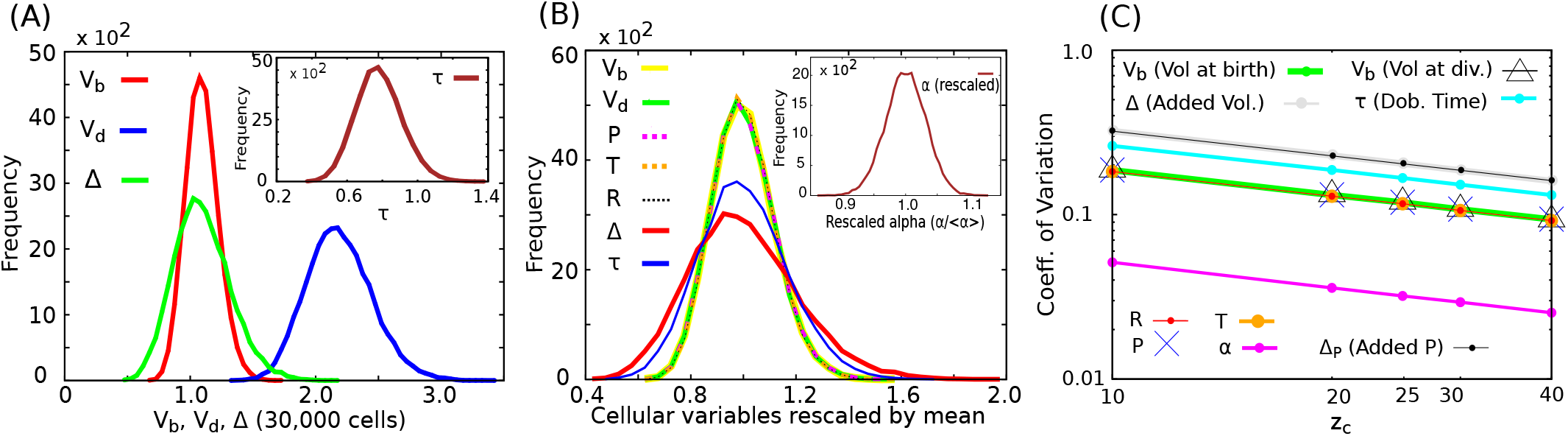
Distribution of volume, interdivision time, growth rate and chemical populations in the PTRZ model. This figure shows the frequency distributions of various cellular variables, measured across 30000 cells at the steady-state. Data was compiled from each trajectory in the 30th generation to ensure that it had reached the statistical steady-state. **A** Distributions of volume-at-birth *V_b_*, volume-at-division *V_d_*, added volume Δ ≡ *V_d_* − *V_b_* and (in inset) interdivision time *τ*. **B** Frequency distributions of various quantities rescaled by their respective means. The *x*-axis for the curve denoted by *V_b_* in the legend stands for *V_b_*/〈*V_b_*〉, and similarly for the other curves. The distributions of rescaled *P_b_, T_b_, R_b_, V_b_* and *V_d_* are essentially indistinguishable. The distribution of rescaled *α* is shown in the inset of **B. C** CV of Δ, *τ, V_b_, V_d_, α, P_b_, T_b_, R_b_* and Δ_*P*_ ≡ *P_d_* − *P_b_* as functions of *Z_c_* on a log-log plot. The slope of the linear fit is consistent with CV ∝ *N*^−1/2^ for all the quantities. The vertical separation between the lines gives the ratios of the CV of various quantities and they are consistent with what is expected when the adder property holds, Eq. (6). Parameter values are as in Fig. 1 except in **C** where *Z_c_* takes a range of values.

The first three relations are expected to hold if the cell volume grows exponentially with a small variation in the growth rate and the system satisfies the adder property [11]. The last equality relates the fluctuations to the parameters *Z_c_* and *Z_r_* of the present model. This connects the CV of a macroscopic quantity Δ to a potentially microscopic quantity *N*, the change in the population of the *Z* molecule from birth to division. The CV of *P_b_, P_d_* and Δ_*P*_ ≡ *P_d_* − *P_b_* also satisfy (6), as do the corresponding quantities for *T* and *R*.

Using the simulation data of the 30000 stochastic cells, we determine the correlations between the cell size at birth (*V_b_*) and division (*V_d_*), the added volume (Δ) and the inter-division time (*τ*) in the steady-state (Fig. 4). The correlations show that the model exhibits the adder property, because the mean added volume Δ between birth and division is independent of the birth size. These are the kinds of correlations observed in experiments with *E. coli* [10, 11]. Taheri-Araghi et al [11] have also measured the distribution of Δ for a fixed *V_b_* at different values of *V_b_* (binned) for *E. coli*. They find that not just the mean of Δ but the entire distribution to be independent of *V_b_*, which is a strong version of the adder property. For the PTRZ model in Fig. S2 we exhibit this ‘conditional’ distribution of Δ for different fixed values of *V_b_* (in a bin). The distributions collapse onto each other, showing that the model exhibits this property.

**FIG 4.**
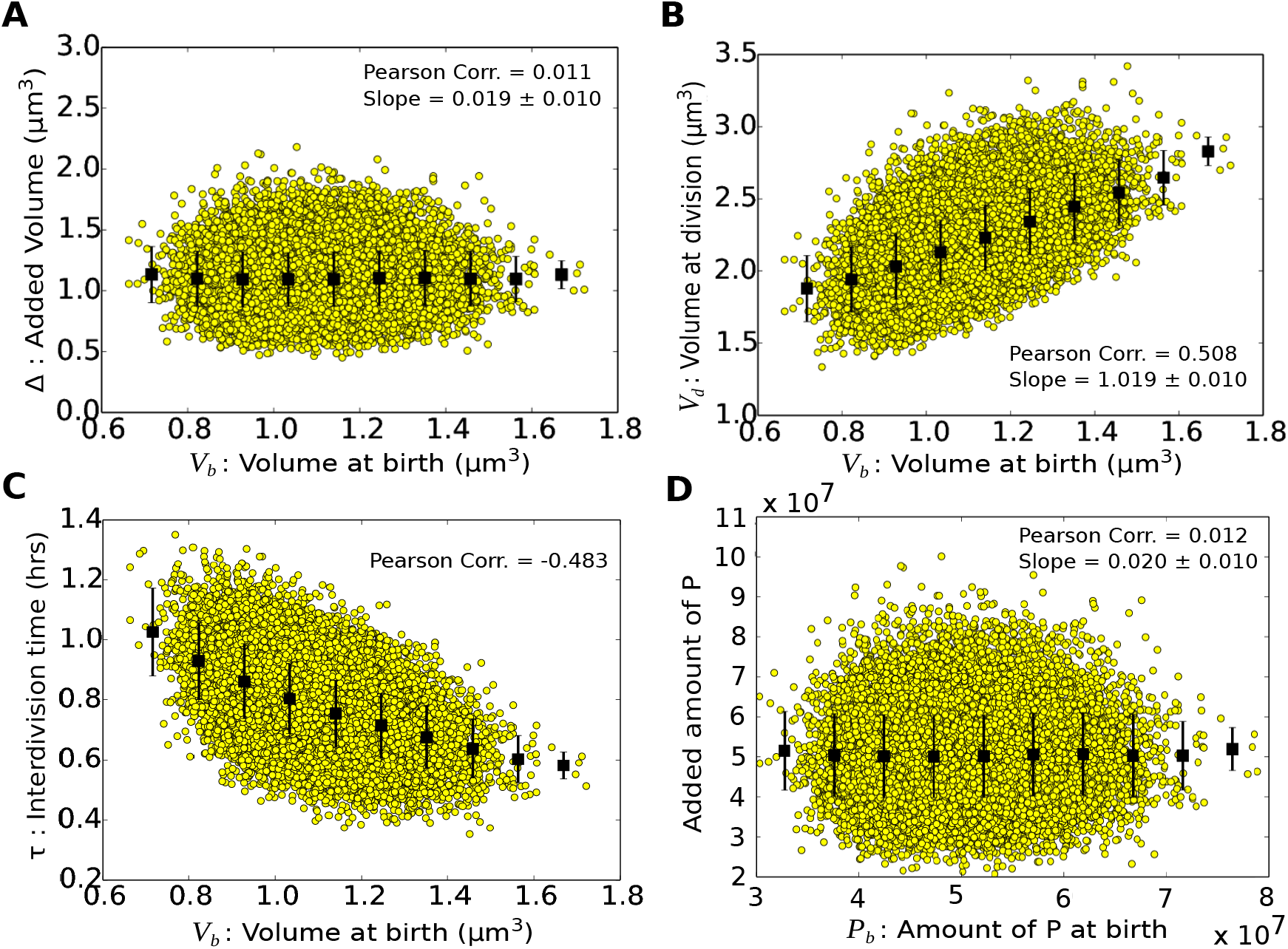
Dynamics of the PTRZ cell model displays the ‘adder’ property for cell size and intracellular chemical populations. Each data point in the scatter plots refers to one trajectory out of 30000. The two axes show two cellular properties pertaining to the 30th generation. E.g., in **A** the *x*-axis shows the volume of a cell at birth and the *y*-axis the added volume of the cell in the same generation. The black squares and vertical lines show the mean and s.d. of Δ within a bin of *V_b_*. The lack of correlation between Δ and *V_b_* is evidence of the ‘adder’ property. The slope (± standard error) of the line of binned data is 0.019 ± 0.010 (we expect zero slope for adder). **B** The volume of the cell at division, *V_d_*, is positively correlated with the birth volume, with slope 1.019 ± 0.010 for the curve of *V_d_* averaged over bins of *V_b_* (for the adder, expect slope unity). **C** The interdivision time *τ* is negatively correlated with the birth volume, with the correlation coefficient between the two variables rescaled by their respective means being -0.483 (expect -0.5 for adder). **D** The increment in *P* from birth to division is uncorrelated with *P_b_*, showing that the chemical population *P* also exhibits the adder property.

##### Adder property is independent of *Z_r_*

We remark that in the present model the adder property of cell volume is observed for the entire range of the reset parameter *Z_r_*, 0 ≤ *Z_r_* ≤ *Z_c_*. This is also shown analytically in section IV D.

#### 3. Intracellular chemical populations: Distribution, adder property and dependence of CV on the mean population

Each population at birth *P_b_, T_b_, R_b_* at birth has its own mean and CV, but the populations rescaled by their respective means have the same distribution which matches the rescaled *V_b_* and *V_d_* distributions (Fig. 3B). This is because the *P,T,R* are all strongly correlated with each other (see Fig. S3), with their ratios dominated by the deterministic dynamics. By virtue of (3) they are also strongly correlated to *V*. We remark that strong correlations between high copy number proteins are also observed experimentally [31]. Further, universal fluctuations in protein populations within cells have been reported in [32].

We find that in the PTR model with stochastic chemical dynamics, the chemical populations *P,T* and *R*, like the volume, also exhibit the adder property (see Fig. 4D for *P*, and Fig. S4 for *T, R*). The reason for these populations showing the same distribution and the adder property is discussed later in section IV D where we give an analytical derivation. We also later identify circumstances wherein other chemical populations in the cell depart from the adder.

##### The crossover behaviour of protein number fluctuations

A slight extension of the present model explains the crossover behaviour of the CV of protein levels as a function of their mean levels observed in [31]. To see this we introduce another variable *Q* in the *X* sector, with 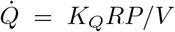. This has the same form as the production of other proteins in (2).

The system now has 5 variables PTRQZ. By dialing Kq we can control the absolute value of *Q* in the steady state. We perform stochastic simulations (with stochasticity in the chemical dynamics of all 5 molecules PTRQZ) using different values of *K_Q_* and determine the mean value of *Q* at birth 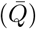 and CV of *Q* for each. The result (Fig. 5) shows a universal 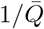 behaviour for small 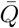 and a plateau for large 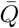, as observed in [31].

**FIG 5.**
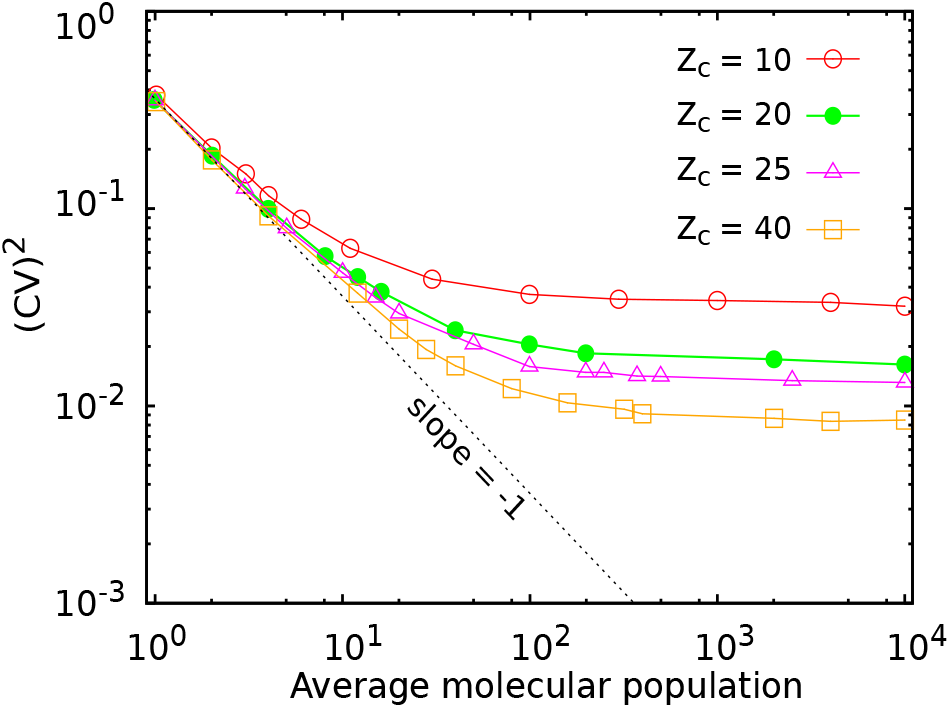
Dependence of CV of a protein population on its mean value. Simulation results of the PTRZ model with an additional protein population *Q* as discussed in section III B 3 are shown. A crossover from a 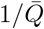 behaviour of (CV) to a constant value is seen at 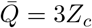 and (CV)^2^ = 1/(3*Z_c_*).

##### An intrinsic source of ‘extrinsic’ noise

In the model there is a simple explanation of the above behaviour. The (CV)^2^ due to intrinsic fluctuations is ~ 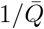 and due to division control is 1/(3*Z_c_*) (from Eq. (6) using *Q* in place of *V_b_*). For small 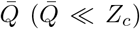 the intrinsic fluctuations dominate; for large 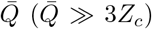 the 1/(3*Z_c_*) term which is independent of 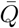 dominates. A protein population has intrinsic fluctuations from stochasticity in production and degrading processes [30, 54]. The variation not explained by intrinsic fluctuations is referred to as ‘extrinsic’ [55]. Stochasticity in the division time has been recognized as a source of extrinsic noise leading to this crossover behaviour [56]. In the present model both the saturating value of (CV)^2^ (≃ 1/(3*Z_c_*)) and the scale of 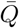 at which the crossover occurs 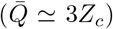 are governed by the division threshold *Z_c_*(or in general by *N* = *Z_c_* − *Z_r_*/2). The so called ‘extrinsic noise’ in the division time is, in the present model, ultimately a consequence of intrinsic fluctuations in the *Z* population, which causes the fluctuations in the first passage time of *Z* reaching its threshold *Z_c_* and hence determines a lower bound on cell-to-cell variation of size and of intracellular chemical populations.

### 4. Comparison with experimental distributions

We compare the distributions of certain quantities rescaled by their means as predicted by the model with the experimentally observed distributions for *E. coli* [11] in Fig. 6. As mentioned earlier the distributions of rescaled Δ, *τ, V_d_* predicted by the model are robust and depend only on a single parameter *N* = *Z_c_* − *Z_r_*/2. Figs. 6 ABC show that the experimental data points for all three quantities largely lie between model curves corresponding to *N* = 20 and *N* = 60, when the only source of stochasticity in the model is the intrinsic dynamics of the populations. (*N* = *Z_c_* in Fig. 6 since *Z_r_* =0). This places an experimental constraint on a key parameter of the model, *N*. Note that in the real cell there could be other upstream sources of stochasticity in *Z* production, such as the stochasticity in its mRNA production and degradation. These have not been taken into account in our simple model. They would also contribute to the CV of the first passage time distribution thereby allowing larger values of *N* to be consistent with the data.

**FIG 6.**
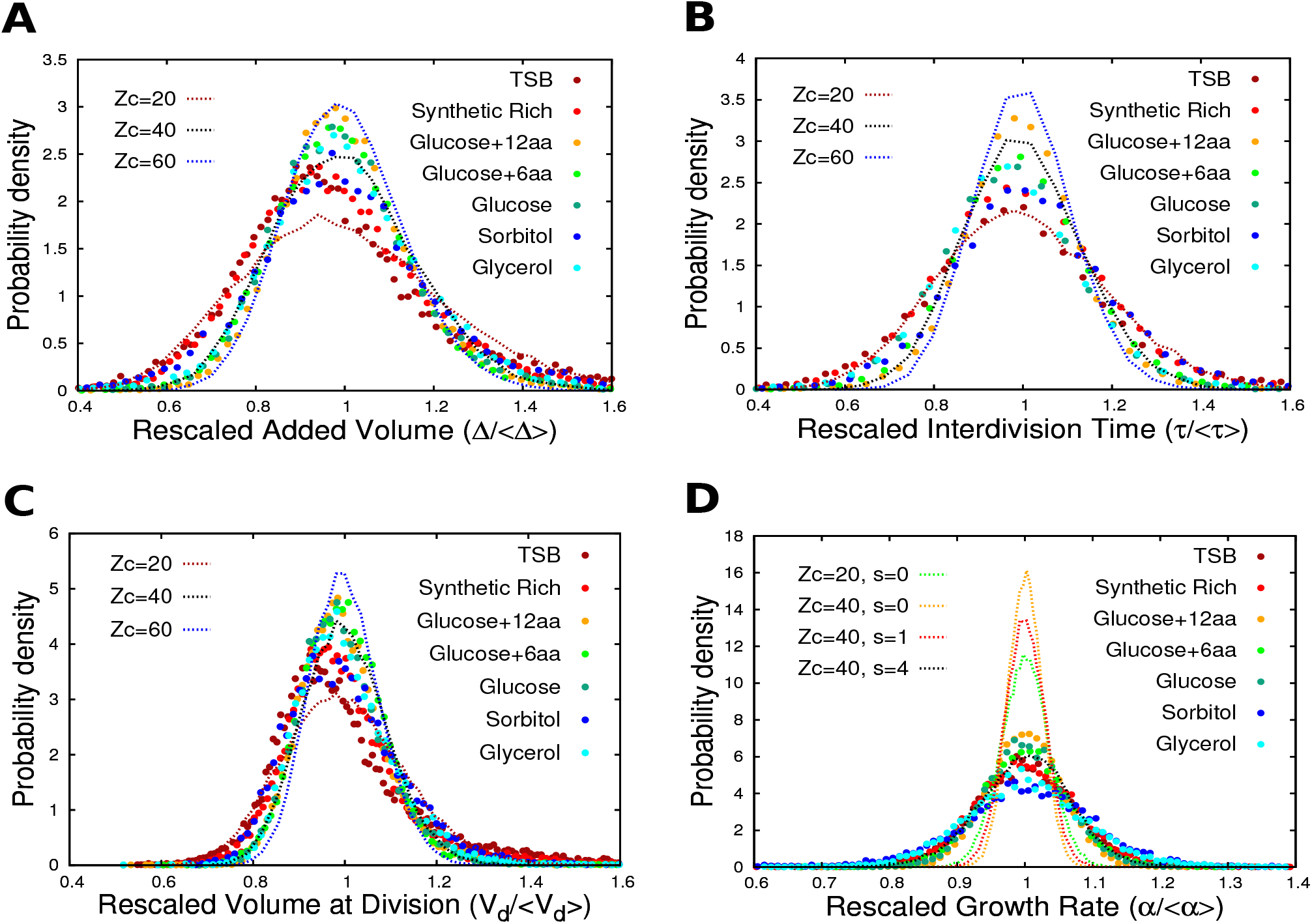
Comparison of steady state distributions predicted by PTRZ model with experimental data. **A,B,C,D** respectively show the distributions of Δ, *τ, V_d_* and *α* rescaled by their means. Experimental values of various quantities for *E. coli* in seven media were obtained from data of [11] to produce the binned histograms. The different media are represented by dots of different colour. Model simulations are done with parameters values as in Fig. 1, except *Z_c_* which takes values 20,40,60. All model histograms (dotted lines) are obtained from an ensemble of 30000 cells. In all simulations the intrinsic stochasticity in the dynamics of all four chemicals P,T,R,Z is present. Partitioning stochasticity is absent in **A,B,C** and in two curves of **D** (*s* = 0; see section IIIB 5 for definition), but is present in two simulations shown in **D** (*s* = 1,4). Threshold stochasticity is absent in all simulations shown (*s*′ = 0).

It is seen in Fig. 6D that in the range *N* ≥ 20 the model predicts a much narrower width for the *α* distribution than is experimentally observed (compare the *Z_c_* = 20, 40; *s* = 0 curves for the model in Fig. 6D with those of the data). While [11] reports a CV ranging from 6-11% depending upon the medium in the experimental data, the model with only intrinsic stochasticity (*s* = 0, see below) predicts a CV of the *α* distribution to be only 3.6% at *N* = 20, 2.5% at *N* = 40 and lower for larger values of *N*. We recall that the *α* distribution in the model is not as universal as the distributions of Δ, *τ, V_d_*, and depends on other model parameters including the parameter values in the PTR sector (Fig. S1). We show below that its width is sensitive to and increases significantly with the inclusion of partitioning stochasticity in the model. In other words, the model allows room to get the experimentally observed width of the *α* distribution also, without destroying its agreement with the other distributions. It is also worth mentioning that there is a significant variation in the CV of the *α* distribution reported by different groups, suggesting that this quantity is sensitive to experimental details and perhaps needs to be measured with a higher degree of control.

### 5. Effect of other sources of stochasticity on the distributions and the adder property

#### Partitioning stochasticity

In addition to the intrinsic stochasticity in the chemical population dynamics, consider a second source of stochasticity, the uneven partitioning of chemicals between the two daughter cells during cell division. Thus at division instead of replacing *X_i_* by *X_i_*/2 we replace it by 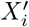 drawn (independently for each *i* and each generation) from a gaussian distribution with mean = *X_i_*/2, standard deviation 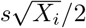, where *s* ≥ 0 is a parameter characterizing the strength of partitioning stochasticity [57] *s* = 1 corresponds to the symmetric binomial partitioning wherein each molecule can independently go to either daughter with equal probability (the gaussian 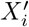 distribution approximates the binomial distribution 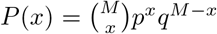 with *M* = *X_i_* and *p* = *q* =1/2). *s*> 1 implies that fluctuations are stronger than the binomial case, a situation that can arise if molecule clusters are partitioned instead of single molecules [57]. The result for the *α* distribution with both sources of stochasticity (intrinsic + partitioning) is shown in Fig. 6D and for the other distributions in Fig. S5. It is seen that when partitioning stochasticity is included (e.g., *Z_c_* = 40, *s* = 4 in Fig. 6D), the model reproduces a width of the *α* distribution comparable to the experimental data (CV of *α* is 6.8% for these parameter values). Fig. S5 shows that the inclusion of partitioning stochasticity has a relatively small effect on the width of Δ, *τ, V_d_* distributions in this regime, and that the adder property continues to hold in the model at these strengths of the partitioning stochasticity.

#### Stochasticity in the threshold value of *Z*

The biochemical mechanism implementing the trigger when *Z* reaches *Z_c_* is expected to have its own stochasticity. Thus the value of *Z* at which division is triggered need not be precisely *Z_c_* in every cell, but could vary from cell to cell and generation to generation around *Z_c_*. We implement this ‘threshold stochasticity’ by drawing the threshold (now denoted 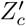) from a gaussian distribution with mean *Z_c_* and standard deviation *s*′, independently for each cell in every generation. The parameter *s*′ ≥ 0 characterizes the strength of the threshold stochasticity. We ran simulations of the PTRZ model with two sources of stochasticity, the intrinsic stochasticity of the chemical dynamics of the populations *P,T,R,Z* and the threshold stochasticity implemented as above. In these simulations the local dynamics of the populations is as before, except that the value of *Z* at which division is triggered, 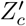, is chosen from the above mentioned gaussian distribution in each generation. Immediately after triggering, *Z* is reset to *Z_r_*, and at division, all populations are halved. We consider two cases (a) *Z_r_* is a fixed number, the same for all generations, and (b) *Z_r_* is a fixed fraction of 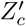 (the reset value of Z after triggering is a fixed fraction of the value of *Z* at triggering).

Threshold stochasticity modifies the distributions of Δ,*τ, V_d_*; however the modification remains small as long as 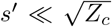 as the dominant contribution comes from intrinsic stochasticity in this regime. When 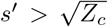, threshold stochasticity is the dominant contributor to these distributions. However, while it modifies these distributions, in case (a) it does not affect the adder property, which survives even when 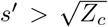. This is shown in Fig. S6A. In fact it can be shown analytically that when threshold stochasticity is the only stochasticity present (the intrinsic stochasticity is turned off), the PTRZ model exhibits the adder property in the steady state, provided that *Z_r_* is independent of 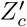. However, when *Z_r_* is correlated with 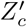, as is true for case (b), we lose the adder property (see Fig. S6B). This is because the added volume from one trigger to the next depends upon the reset value *Z_r_* after the first trigger, while *V_b_* of a generation depends on the value of 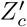 in the previous generation. Thus the added volume becomes correlated with *V_b_* when *Z_r_* is correlated with 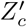.

Unlike partitioning stochasticity, threshold stochasticity does not contribute significantly to the width of the *α* distribution even though it affects the Δ, *τ, V_d_* distributions. In fact, when threshold stochasticity is the only stochasticity present, the latter distributions have a finite CV but the CV of the *α* distribution is zero. The reason for this will be given in section IV E.

Another kind of stochasticity present in the cell is that the position of the septum is not necessarily at the middle of the dividing cell. This directly affects the distribution of *V_b_* and enhances its CV, which is found to be larger than the CV of *V_d_* [11]. The effect of this stochasticity is not considered in the present paper.

## IV. ANALYTICAL DERIVATION OF THE RESULTS OF THE MODEL

In this section we provide explanations of the results of the PTRZ model presented in the previous section. In particular we discuss the exponential growth of populations and *V*, and give analytic expressions for the average interdivision time scale, intracellular concentrations, and the cell size. We also explain the origin of the adder property and the shape of the Δ and *τ* distributions. We explain why the *α* distribution behaves so differently from the Δ, *τ, V_d_* distributions under different types of stochasticity and what factors contribute to it. These results will be generalized to a much wider class of models than the PTRZ model in section V.

### A. The PTR sector, exponential growth and the growth rate

At the deterministic level, all the features of the steady state solution in Fig. 1 can be understood by considering an exponential ansatz for the trajectories of the chemical populations:

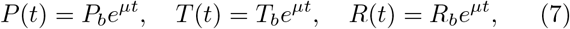

with 4 unknown constants *P_b_,T_b_, R_b_* (representing the populations at birth of the cell in the steady state), and *μ*, the growth rate of the PTR cell. It is easy to see [50] by substituting (7) into Eq. (2) that (7) is a solution of (2) only if the ratios of the populations and *μ* are fixed in terms of the parameters. Specifically,

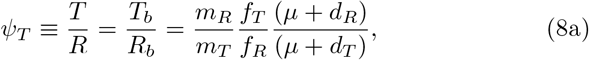

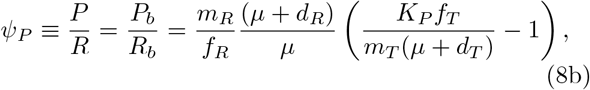

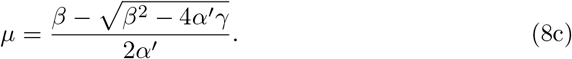

Here *α*′ = 1 − *ϵ*_1_, *β* = *a* + *b* + *ϵ*_2_,*γ* = *ab; a* = *νf_T_* − *d_T_, b* = *ρf_R_* − *d_R_; ν* = *K_P_*/*m_T_,ρ* = *k*/(*m_R_υ_P_*); 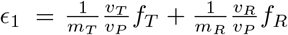, 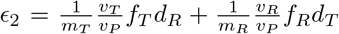. If *P_b_, T_b_, R_b_* satisfy (8a),(8b) and *μ* is given by (8c), then (7) is a solution of (2). That an exponentially growing trajectory is a solution of the nonlinear equations (2) is a consequence of the fact that the right hand sides of (2) are homogeneous degree one functions of the populations: *f_i_*(*βP, βT, βR*) = *βf_i_*(*P, T, R*), *i* = 1, 2, 3. This in turn is a general consequence, as will be discussed later, of the assumption of the mass action chemical kinetics implicit in (2) and the linearity of V in the populations (3).

Not only is (7) a solution of (2), it is a stable attractor of the dynamics. In Fig. 1 we have considered the PTR sector in conjunction with the *Z* sector which truncates the growth of PTR at discrete times (at the point of division Eq. (2) is effectively suspended). However if there is no such truncation and (2) together with (3) is the sole dynamics of PTR, then the exponentially growing solution (7) happens to be an asymptotic attractor of the dynamics (see Fig. S9 for numerical evidence with different initial conditions and parameter sets). Thus, starting from an arbitrary initial condition, eventually the system approaches the exponential trajectory (7) with *μ* given by (8c) and ratios of populations given by (8a) and (8b).

When division control via the *Z* sector is included in the dynamics, we get the behaviour shown in Fig. 1 with a limit cycle attractor. In the growth phase of this attractor (the period after birth and just before division), the populations again grow exponentially following (7), with the exponential growth rate of *P, T, R, V* matching the formula (8c), and the ratios of chemicals in the attractor matching (8a), (8b). We have verified this numerically for diverse initial conditions (with *P*(0),*T*(0),*R*(0) > 0 and 0 ≤ *Z*(0) < *Z_c_*) and diverse parameter sets for which the r.h.s. of (8c) is positive (when the latter is negative there is no exponential growth). The numerical work suggests that given a fixed set of parameters (with a positive r.h.s. of (8c)), for arbitrary physical initial conditions the system always settles down in a limit cycle attractor similar to the one described in Fig. 1 such that the trajectory between birth and division in every cycle is described by (7) and (8) (numerical simulations have been done for non-negative values of the rate constants and other parameters of the model). The positivity of the r.h.s. of (8c) seems to be the only collective requirement on the parameters of the PTRZ model for this kind of dynamical behaviour to arise.

### B. Curve of balanced growth, interdivision time, concentrations

By construction the dynamics of the PTR sector does not depend upon *Z*, except for the fact that at certain discrete times (when *Z* approaches *Z_c_*) the 3 populations are halved. Therefore it is useful to consider the projection of the dynamics onto the 3-dimensional space with coordinates (*P, T, R*). (Since we are dealing with populations, we only consider the positive octant.) Geometrically, the eqns. (8a) and (8b) define a straight line passing through the origin of this 3-dimensional space whose angles with the 3 coordinate axes are fixed by the parameters. We refer to this line in the 3-dimensional space as the ‘curve of balanced growth’ (CBG) for this system. If the initial point of a trajectory lies on this line, the 3 populations grow exponentially according to (7) with the rate *μ* given by (8c), and their ratios remain constant in time and given by (8a)(8b). Since the division process halves the 3 populations, they remain on the CBG after division. Thus the CBG is an invariant manifold of the deterministic dynamics. Since numerically we find that starting from arbitrary initial values the ratios approach those given by (8a), (8b), this means that the stable attractor of the dynamics lies on the CBG. In fact the steady state is a limit cycle lying on the CBG characterized by repeated rounds of exponential growth of populations from birth to division with growth rate *μ* until the populations double, followed by halving of the populations. The interdivision time scale on this limit cycle is therefore given by

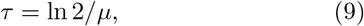

with *μ* given by (8c). Note that the concentrations of three chemicals, given by [*P*] ≡ *P/V*, [*T*] *T/V* ≡ *T/V* and [*R*] ≡ *R/V* are the same everywhere on the CBG and hence constant on the limit cycle. This is because *V* is a linear function of the populations (3). Thus, for example, *V* = *R*(*υ_P_ψ_P_* + *υ_T_ψ_T_*+*υ_R_*); hence *R/V* is completely expressed in terms of the ratios *ψ_P_* and which are constant on the CBG. Thus the growth rate or interdivision time scale and all intensive quantities pertaining to the PTR sector in the steady state of the deterministic dynamics are completely determined by the parameters of the PTR sector of the model.

### C. The *Z* sector and cell size

The *Z* sector determines the absolute size scale of the cell by fixing one extensive quantity pertaining to the PTR sector. In the deterministic steady state since *P, T, R* satisfy (7), we can write (5) as

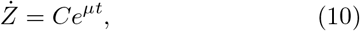

where 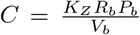. *C* is an extensive quantity of the PTR sector (homogeneous degree one in the populations) and can be written in terms of *V_b_* and the intensive quantities:

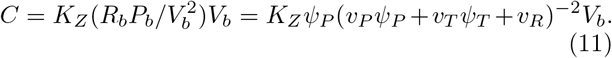

The requirement that in the steady state *Z* must also complete its cycle in the doubling time *τ* fixes *C* and hence the size of the cell. (10) has the solution

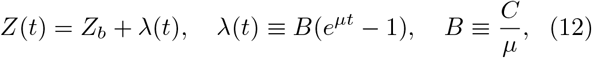

where *Z_b_* is the value of *Z* at *t* = 0. Note that unlike *P, T* and *R, Z* does not increase exponentially because it is reset from *Z_c_* to a value *Z_r_* ≠ *Z_c_*. If there is no resetting (*Z_r_* = *Z_c_*), then (12) implies that *Z* also increases exponentially.

When *τ*_1_ = 0, then *Z* equals *Z_c_* at *t* = *τ*, and *Z_b_* = *Z_r_*/2. Substituting this in (12) gives *B* = *N* ≡ *Z_c_* − *Z_r_*/2. This fixes the absolute size scale of the cell and the absolute populations. In particular we get *R_b_* = *N_μ_K_Z_ψ_P_*/(*υ_P_ψ_P_* + *υ_T_ψ_P_ + υ_R_*), from which *P_b_,T_b_* can be obtained by multiplying by *ψ_P_,ψ_T_*. Further, using (11) we get

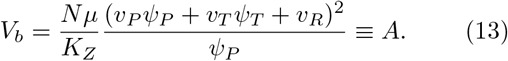

This is an explicit formula for the cell size at birth in terms of the model parameters (*ψ_T_* and *ψ_P_* being given by (8a),(8b)). The numerical values obtained in deterministic simulations (e.g., Fig 1) agree with this formula (as do the absolute populations).

When *τ* > *τ*_1_ > 0, instead of *B* = *N* we get *B* = *Ne*^*μτ*_1_^ (see Supplementary section S2A for the derivation). Then it follows that

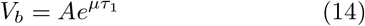

where *A* is defined in (13). This expression contains the exponential factor *e*^*μτ*_1_^ = *e*^*μ*(*C+D*)^ obtained by Donachie [58]. The *e*^*μτ*_1_^ factor in the expression for *V_b_* arises because of the exponential dependence of *Ż* on time, Eq. (10), which is a consequence of the exponential growth of the *X* sector populations in the steady state. The latter, in turn, is a consequence of the homogeneous degree one nature of the functions *f_i_* defining the *X* sector dynamics which we argue later is quite universal and not restricted to the PTR model.

The PTRZ model has been defined above for *τ* > *τ*_1_. To extend it for higher growth rates one needs multiple origins of replication [59]. Adapting the work of [36] to the present model it can be shown that the formula (14) holds for higher growth rates as well. Details will be presented elsewhere.

The Schaechter et al growth law of cell size [40], a strong version of which has been established in [60], states that the average cell volume in different growth conditions depends exponentially on the growth rate *μ*. Note that while Eq. (14) contains the exponential factor *e*^*μτ*_1_^ it cannot be construed as equivalent to this growth law. This because (14) has, in addition to the exponential factor *e*^*μτ*_1_^, the prefactor A which itself has a complicated *μ* dependence. This factor depends upon the ratios of various chemicals in the steady state (which, in turn, depend upon *μ*; see (8)) and the contribution of each chemical to the volume of the cell (the constants *υ_i_*). In general the prefactor *A* (unlike the *e*^*μτ*_1_^ factor which is much more universal) depends upon the details of the *X* and *Z* sectors - the actual form of the functions *f_i_, h*, regulatory mechanisms acting in the X sector (which are contained in the functions *f_i_*), parameter values, etc. A more general expression for A in terms of the functions *f_i_, h* and *υ_i_* is given later, Eqn. (34). However, comparing the theoretical prediction (14) or (34) with experiments, in particular reproducing the growth law [40, 60], is a task for the future.

### D. Stochastic dynamics and the adder property

The full stochastic dynamics of the PTRZ model whose numerical results were presented earlier is difficult to treat analytically. However, we can make some approximations to obtain partial results. As observed numerically in the statistical steady state the growth of PTR sector populations in a given generation could be approximated by an exponential fit (Fig. 2B). Moreover since the actual numbers of *P, T* and *R* were large, their relative fluctuations around their average trajectories were small. Thus as an approximation we ignore the stochasticity in the chemical dynamics of *P, T, R* and only consider the stochasticity in *Z*. For simplicity we assume that in the steady state the PTR dynamics is deterministic and lies on the curve of balanced growth discussed earlier. Thus *P*(*t*), *T*(*t*), *R*(*t*) are assumed to be given by (7), their population ratios and the growth rate being constant and fixed by (8). However, due to the fluctuation in the time taken by *Z* to reach *Z_c_*, the exponential factor by which they grow varies from generation to generation and in each generation they start from a different point on the CBG at birth (i.e., the absolute scale of *P_b_, T_b_, R_b_* varies from generation to generation). Clearly this approximation assumes that there is no fluctuation in the growth rate *α* and hence we cannot hope to obtain the *α* distribution from this approach (*α*, defined as the slope of ln *V* vs *t*, equals *μ* in this approximation, whose value is given by the r.h.s. of (8c)). However, we can obtain analytic expressions for the *τ* and Δ distributions.

#### 1. Derivation of the adder property and distributions of τ and Δ

In order to get the probability distribution of the interdivision time, we need to consider the stochastic version of the differential equation (1b), or more specifically, (5). As a consequence of the assumptions in the previous paragraph we can use (7), hence (5) reduces to (10). This means that the probability that *Z* increases by unity in the small time interval (*t, t* + *δt*) is given by *Żδt* where *Ż* is given by (10), and the probability that it remains unchanged is 1 − *Żδt*. A given trajectory starts with fixed values of *P, T, R, V, Z* at birth, denoted *P_b_, T_b_, R_b_, V_b_, Z_b_* and hence a fixed value of 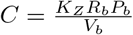. These values can change from generation to generation, but since in the present approximation *P_b_, T_b_, R_b_* are assumed to always lie on the CBG, the ratio *C*/*V_b_* is constant for all generations and given by (11). This justifies a crucial assumption, namely, that *C* ∝ *V_b_* or equivalently *Ż* ∝ *V_b_e^μt^*, made in [37] where the first passage time problem of a division triggering chemical based on (10) has been discussed. Consider the ensemble of trajectories with a fixed value of *P_b_, T_b_, R_b_, Z_b_*, and hence fixed *V_b_* and *C*. Since the r.h.s. of (10) is a fixed function of time, the stochastic process in *Z* is the 1-dimensional inhomogeneous Poisson process (if the r.h.s. had been independent of t it would have been the standard Poisson process). Given *Z* = *Z_b_* at *t* = 0, the probability that the first passage time for it to reach *Z_c_* is between *τ* and *τ* + *dτ* is given by 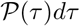, with (see Supplementary section S2 B for a derivation)

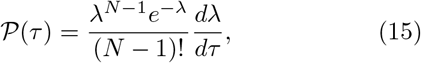

where *λ* = *λ*(*τ*) = *B*(*e^μτ^* − 1) and *N* = *Z_c_* − *Z_b_* = *Z_c_* − *Z_r_*/2.

In a given trajectory of the ensemble, the volume increment between birth and division is given by Δ = *V_b_*(*e^μτ^* − 1) = (*V_b_/B*)*λ*(*τ*). This being proportional to *λ*, the probability 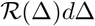 that a given trajectory in this ensemble has a volume increment in the range Δ to Δ + *d*Δ is obtained from (15):

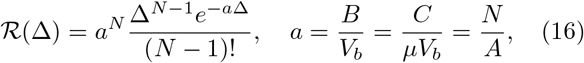

where *A* is given by (13). From this it follows that

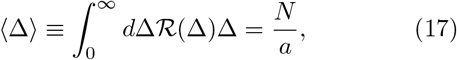

which is the same as the deterministic value of *V_b_* (13). The distribution of rescaled Δ, *u* ≡ Δ/〈Δ〉 is thus a Gamma distribution given by

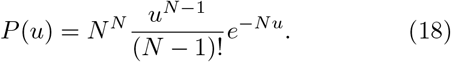

The adder property follows from (16). Note that *N* = *Z_c_* − *Z_r_*/2 is a constant parameter of the model that does not change from generation to generation as long as *Z_c_, Z_r_* do not change from generation to generation. The only other parameter that determines the shape of 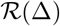 in (16) is a which is independent of the value of *V_b_* in any generation. The latter is the case because on the CBG, where (10) holds, a is a constant irrespective of the scale of *V_b_* (*a* = *C*/(*μV_b_*) with *C* given by (11). Thus *a* depends only on the *ratios* of populations of the PTR sector, and on the CBG these ratios are the same irrespective of the scale. This proves the strong version of the adder property: not just the mean but the entire distribution of Δ is independent of *V_b_*. The distribution of *u* only depends upon *N* and on no other parameter of the PTRZ model. This explains the extraordinary robustness of the *u* distribution mentioned in Section IIIB 2. From (18) it follows that CV of 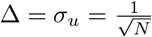, which derives the last equality in (6).

The above distributions and the adder property were obtained in [37] from the assumptions that the cell volume grows exponentially with time (*V*(*t*) = *V_b_e^μt^*) and that the rate of growth of the time-keeper protein *Z* is proportional to *V*(*t*). The first assumption is equivalent to a linear model for cell size growth, 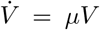. The present work obtains the exponential dependence of *V* on *t* from a non-linear dynamical model of the cell whose volume is defined in terms of the three chemical populations. As discussed above it also justifies the second assumption (*Ż* ∝ *V*). In other words, in the present work Eq. (10) with *C* ∝ *V_b_* is not an assumption but a consequence of more basic dynamics, wherein the cell is attracted to the CBG. We show later in section V C that this holds for a much larger class of models in which the *X* sector has *n* chemicals where the functions *f_i_* and *h* in Eq. (1) are nonlinear functions arising from mass action kinetics. Thus the present work generalizes the results of [37] to a very large class of nonlinear models.

We note that the present model implies that the populations of the *X* sector also satisfy the adder property. Their increments between birth and division, Δ_*i*_ ≡ *X_id_* − *X_ib_*, have the same distribution as Δ given by (16), with *a* = *B/X_b_*. This follows from the fact that like *V*, the *X_i_* also grow exponentially, hence Δ_*i*_ = *X_b_*(*e^μτ^* − 1) = (*X_b_/B*)*λ*(*τ*).

In the above derivation of the Δ and *τ* distributions and the adder property it has been assumed that *P, T, R* have no stochasticity and lie on the CBG. When stochasticity in *P, T, R* is included, they no longer lie on the CBG. In fact in a given generation the best fit value of the growth rate is not the same for all three populations, hence (5) does not reduce to (10). Nevertheless the analytically derived distributions (18) and (15) compare well with our numerical simulations of PTRZ model where stochasticity in *P, T, R* is included; see Fig. 7A,B. Further as discussed in section III B 2 and shown in Fig. 4 the adder property also appears. This suggests that the approximation of treating *P, T, R* as deterministic and on the CBG is a good approximation to the full stochastic dynamics.

**FIG 7.**
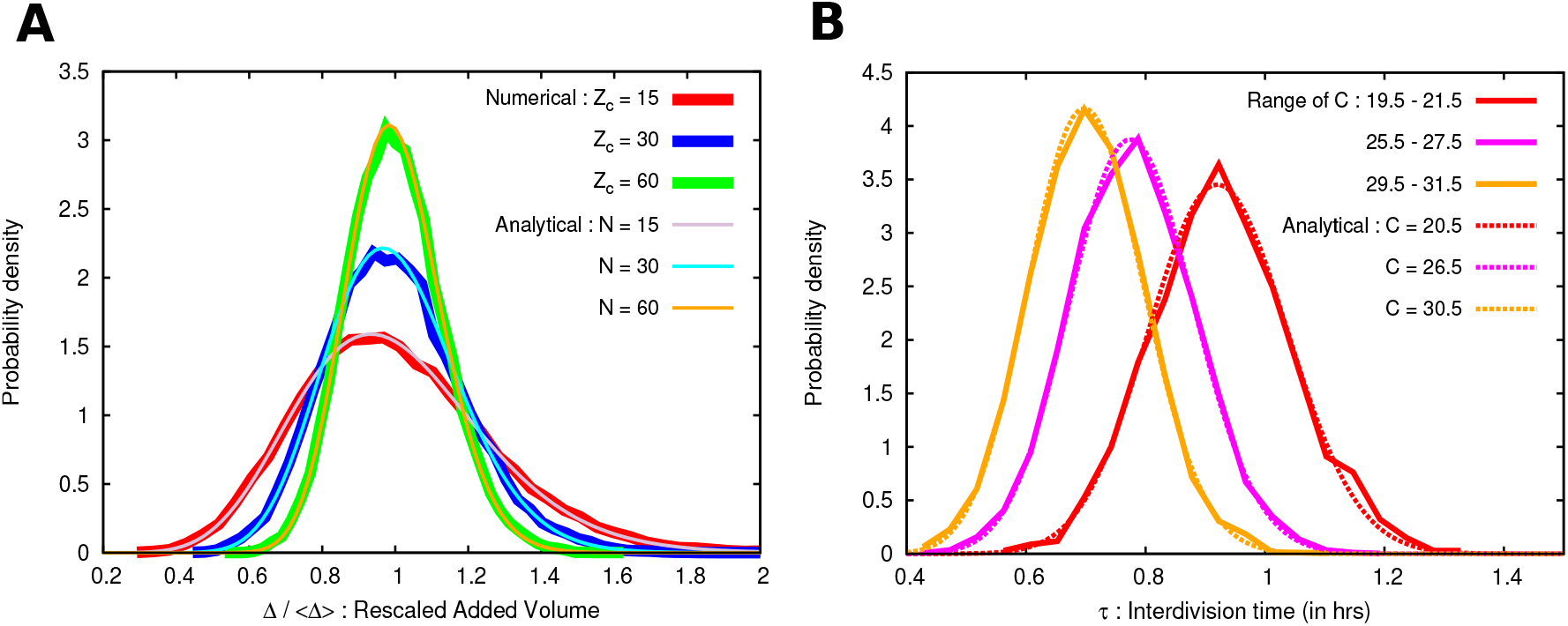
Analytic Δ,*τ* distributions Eqs. (18) and (15) agree with simulations of PTRZ model. **A** Rescaled Δ distribution (18) compared with numerical simulations of PTRZ model for different values of N. Data from all bins of Vb has been pooled together in the numerical curves, since the Δ distribution is independent of *V_b_* (see Fig. S2). **B** Conditional *τ* distribution obtained from numerical simulations of PTRZ model compared with Eq. (15). Since the analytical distribution is conditional on the value of *C*, for comparison the numerical data was binned in ranges of *C* indicated in the legend. The derivation of Eqs. (18) and (15) assumes stochasticity in *Z* dynamics only; *P, T, R* are assumed to lie on the CBG and obey deterministic dynamics. In the PTRZ simulations there is stochasticity in the chemical dynamics of *Z* as well as *P, T* and *R*. The parameters for the PTRZ model are the same as in Fig. 1, except that *Z_c_* = 15, 30, 60 in **A** and *Z_c_* = 30 in **B**.

#### 2. When does the adder property arise?

The adder property and the Δ and *τ* distributions have been derived above from three assumptions: **(i)** in a given generation the cell volume obeys *V*(*t*) = *V_b_e^μt^*, where *V_b_* is the volume at birth in that generation (taken to be at *t* = 0), **(ii)** a molecular population *Z* which starts at some value *Z_b_* triggers division at time *τ* when it reaches a threshold value *Z_c_*, where *Z_c_* − *Z_b_* is a positive constant (the same for all generations), and **(iii)** the dynamics of *Z* between birth and division is the stochastic version of (10) in which the constant *C* is such that *C/V_b_* is a constant for all generations. In fact the same can be derived from weaker assumptions that do not require *V* and *Z* to be exponential functions of time. Consider the case where *Z* and some quantity *Y* have their rates of increase between birth and division to be proportional to each other:

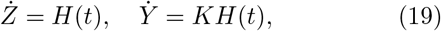

where the function *H*(*t*) and the constant *K* could depend upon the generation. If now we make the *Z* dynamics the stochastic version of *Ż* = *H*(*t*), while retaining the deterministic dynamics of *Y, Ẏ* = *KH*(*t*) and also retaining assumption **(ii)** above, the distribution (15) still arises, with 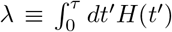 (see Supplementary section S2B). In other words (15) gives the conditional distribution of interdivision time, valid for those generations in which the function *H*(*t*) is the same. Now 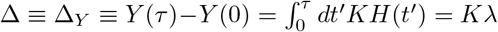, hence the distribution of Δ is again given by (16) with *a* = 1/*K*. This distribution is independent of the functional form of *H*(*t*), and only depends upon *K*. In other words *H*(*t*) appearing in (19) does not have to be an exponential function of time as in (10). Thus if *K* is independent of *Y_b_* = *Y*(0), then so is 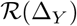, and therefore Y displays the adder property. This argument shows that not only the adder property for *Y* but also the shape of the distribution (16) for Δ_*Y*_ arises when these assumptions hold (namely, (ii), (19) with *K* independent of the generation, and the stochastic dynamics of *Z*).

A biochemical scenario in which Eq. (19) can arise was presented by Sompayrac and Maaloe [33] and emphasized recently in the context of the adder property [35, 36]. In that scenario the *Z* molecule is on the same operon as another molecule *A* whose concentration is held constant in the cell through regulatory mechanisms (autorepression). This implies that the increase in cell volume is proportional to the number of molecules of *Z* produced, thereby realizing Eq. (19) with *H*(*t*) an unspecified function of *t*. Then together with assumption **(ii)** and the stochastic dynamics of *Z*, the adder property of cell volume and the distributions (15) and (18) follow as argued above.

The present model presents an alternative mechanism for realizing Eq. (19) which does *not* require a regulatory mechanism. Here this property arises because the population dynamics has an attractor lying on the CBG in which both *Ż* and *V* are proportional to *e^μt^*. We argue later that the latter is a very generic property of chemical dynamics in self-expanding containers. As a consequence it predicts that the adder property of the volume should be accompanied by the same for intracellular populations (the variable *Y* could be *V* or any of the *X_i_*).

##### Cooperativity in the *Z* dynamics does not spoil the adder property

We now consider the generalization in which the function *h*(**X**, *Z*) in (1b) has a nontrivial *Z* dependence. In particular we consider the generalization of (5):

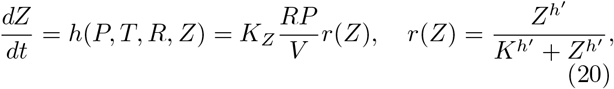

where *K* and *h*′ are constants. *r*(*Z*) represents a positive auto-regulation of *Z* production, *h*′ being a Hill coefficient measuring the strength of cooperativity in the *Z* dynamics. Since *Z* does not appear in the functions *f_i_*, in the deterministic dynamics the PTR variables converge to same ratios as before (the CBG is unaffected by this change) and the growth rate *μ* in the steady state is also the same and given by (8). Under the approximation that PTR sector is treated deterministically (discussed at the beginning of IVD), Eq. (10) is therefore replaced by

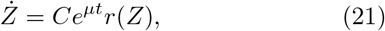

where *C* is given by (11) as before. In order to determine the first passage time distribution and the added volume distribution we now need to consider the stochastic version of (21), where the r.h.s. has a nontrivial *Z* dependence. By considering the master equation following from (21) we show analytically in the Supplementary section S2 C that the presence of a nontrivial auto-regulation changes the shape of the Δ distribution (which is no longer given by (16)), but the adder property *remains intact*. This is also borne out by numerical simulations shown in Figs. S7A and S7B, conducted on the PTRZ model with intrinsic stochasticity in all four population variables and auto-regulation implemented in the *Z* sector via *r*(*Z*). Both positive and negative auto-regulation of *Z* preserve the adder property.

However, the adder property is *not* preserved if the regulatory function *r* is a function of the concentration of *Z*, [*Z*] ≡ *Z/V*, instead of the population *Z* (see Figs. S7C, S7D and text in Supplementary section S2 C).

##### Departure from adder

After the brief excursion in the last two paragraphs to the case where *h* depends non-trivially on *Z*, we return to case where h is independent of *Z* (where the *Z* dynamics is given by (5) and the distributions (15), (16) hold along with the adder property. Since the Δ distribution (16) depends upon *N*, one way of losing the adder property is to have *N* correlated with *V_b_*. The part of assumption (**ii**) that *Z_c_* − *Z_b_* is the same in all generations ensures that *N* is not correlated with *V_b_*. A weaker assumption that *N* varies from generation to generation but is uncorrelated with Vb would still give the adder property, though the distribution for Δ would in general change and depend upon the distribution of *N*. An example of this is given in section III B 5 where threshold stochasticity is considered. We discussed two cases: (a) Where *Z_c_* varies randomly from generation to generation but *Z_b_* = *Z_r_*/2 is fixed; this corresponds to the weaker assumption **(ii)** that preserves the adder property (see Fig. S6A) because *N* is still uncorrelated with *V_b_*. (b) *Z_c_* varies randomly from generation to generation and *Z_r_* of a generation is correlated with the *Z_c_* of that generation; this spoils the adder property (see Fig. S6B) because this correlates the *Z_b_* (and hence *N*) of a given generation with the *V_b_* of the same generation. Since the triggering event at the threshold and the degradation of *Z* must be implemented by biochemical mechanisms, it is possible to imagine both kinds of scenarios.

##### Departure from the adder property for an intracellular chemical population while cell size exhibits the adder property

Up to now we have seen that in the PTRZ model, the adder property for *V* is accompanied by the adder property for the chemical populations *P*, *T*, *R*. However in general it is not necessary that all molecules in the cell exhibit the adder property when the cell size does. We now discuss an example where *P*, *T*, *R* and *V* exhibit the adder property while another molecule *Q* in the cell does not. Consider the PTRZ model defined by Eqs. (2), (3), (5) augmented with another molecule *Q* in the *X* sector whose population dynamics is given by

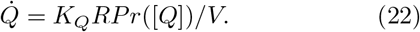

Here *r*([*Q*]) is a function of the concentration of *Q*, [*Q*] ≡ *Q*/*V*, that represents auto-regulation of *Q* production. E.g., *r*([*Q*]) = [*Q*]*^h′^*/(*K^h′^* + [*Q*]*^h′^*) represents auto-enhancement and *r*([*Q*]) = *K^h′^*/(*K^h′^* + [*Q*]*^h′^*) represents auto-repression (with *h′* > 0 being a Hill coefficient, and *K* being another constant). In the model we allow two sources of stochasticity: (i) the intrinsic stochasticity in the chemical dynamics of the molecules *P*, *T*, *R*, *Z*, *Q* and (ii) the partitioning stochasticity at division as discussed in section III B 5. Fig. S8 displays a simulation where V exhibits the adder property but *Q* does not. It is seen that the increment in *Q*, Δ*Q* ≡ *Q_d_* − *Q_b_* (where *Q_b_* and *Q_d_*, are, respectively, the population values at birth and division) increases with Qb in the case of autoenhancement and decreases with Qb in the case of autorepression. This behaviour occurs because partitioning stochasticity causes a departure from the curve of balanced growth as discussed in the next subsection.

### E. Origin of the α distribution

As is evident from the above, under the hypothesis that the first passage time of a molecule to reach a threshold controls cell division, the distributions of τ, Δ and V_d_ are closely related and their CVs are governed by *N* or equivalently *Z_c_* as described by (6). One can now ask for what controls the distribution of α in this setting. Note that *α* is obtained for a given trajectory by fitting the observed trajectory of cell volume to the exponential form *V* (*t*) ~ *e*^*αt*^. Thus *α* in any generation is the slope of the best fit straight line to the ln *V* vs *t* curve from birth to division. In the analysis of the previous subsection where analytic expressions for the *τ* and Δ distributions have been derived, the *X* sector populations *P*, *T*, *R* are assumed to obey their deterministic dynamics and lie on their CBG. On the CBG the deterministic growth trajectory of *P*, *T*, *R* is given by (7), hence the value of *α* is fixed and equal to *μ* given by (8c). Thus the analysis of the previous subsection *assumes* that the *α* distribution has zero width and is therefore just the Dirac delta function *δ*(*α* − μ). That the analysis produces nontrivial distributions of Δ, *τ*, *V_d_* while assuming that the *α* distribution has zero width suggests that any explanation of the origin of a non-trivial *α* distribution must invoke phenomena not included in that analysis.

In our numerical simulations of the PTRZ model we included the stochastic fluctuations of the *P*, *T*, *R* populations. There we found the *α* distribution to have a finite width (Fig. 6D), although smaller than the one experimentally observed in [11]. We noticed that partitioning stochasticity increased the width of the *α* distribution to reach the width observed in [11] as the strength s of partitioning stochasticity was increased. However, threshold stochasticity did not contribute to *α* width. Furthermore while the rescaled Δ, *τ*, *V*_*d*_ distributions depended only on the parameter *N*, the rescaled *α* distribution was found to be dependent on the other parameters of the PTRZ model as well (Fig. S1). These observations suggest that (a) intrinsic stochasticity, and (b) departure from the CBG resulting in a mixing of a pure exponential (given by rate *μ*) with other functions of time contribute to the width in *α* in these models.

The role of intrinsic stochasticity in producing a nonzero width is straightforward. A population variable that is a smooth exponential function of time in a deterministic simulation would be a jagged function when intrinsic stochasticity is switched on, to which an exponential fit would give a different value of growth rate in different trials. The volume being a linear combination of chemical populations will consequently also have a variation in *α* in different generations.

To explain how the departure from the CBG is an independent source of variation of *α* we consider two models in the same broad class defined in section II but linear and much simpler than PTRZ, one containing only one population variable in the *X* sector (the XZ model) and the other containing two (the XYZ model). These are defined by:

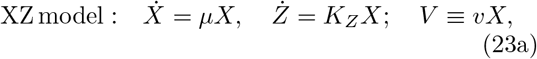

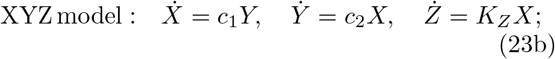

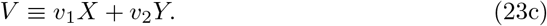

Here *μ*, *K_Z_*, *v*, *c*_1_, *c*_2_, *v*_1_, *v*_2_ are constants. When these models are simulated with only intrinsic stochasticity in the chemical dynamics (partitioning stochasticity and threshold stochasticity absent) they produce the same rescaled Δ, *τ*, *V_d_* distributions as the PTRZ model and their own specific nonzero width *α* distributions (see Fig. S1). This is in keeping with the discussion above. However, now consider simulations of both these models treating the population dynamics of all chemicals deterministically and implementing only partitioning stochasticity with some strength s as described in section III B 5 (intrinsic stochasticity and threshold stochasticity are absent). The resulting distributions are shown in Fig. 8. The Δ, *τ*, *V_d_* distributions are different from what we get when only intrinsic stochasticity is present and all three have nonzero width for both models. But the *α* distribution has zero width for the XZ model and a nonzero width for the XYZ model. This difference between the two models shows that departure from the CBG contributes to the *α* distribution. For, in the XZ model the phase space of the *X* sector coincides with the CBG (both are 1-dimensional) and there can be no departure from the CBG: any initial condition *X*(0) leads to the trajectory *X*(*t*) = *X*(0)*e^μt^*, which always leads to *α* = *μ*. Whereas in the XYZ model the XY sector phase space is 2-dimensional and for a general initial condition (*X*(0), *Y*(0)) the trajectory of the system is a linear superposition of two exponentials: *X*(*t*) = *ae^μt^* + *be^−μt^*, *Y*(*t*) = *α′e^μt^* + *b′e′^μt^*, where *μ* = (*c*_1_*c*_2_)^1/2^ and *a*, *b*, *a′*, *b′* are linear combinations of *X*(0), *Y*(0). Thus *V*(*t*) is also in general a superposition of two exponentials and fitting it to a single exponential will yield a value of *α* that will depend upon *X*(0), *Y*(0). In this linear example the dynamics is governed by the matrix 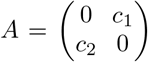, which has two eigenvalues ±*μ*. The eigenvector corresponding to +*μ* is the attractor of the dynamics and is the CBG (it is the line in the XY plane which passes through the origin and has the slope *Y*/*X* = (*c*_2_/*c*_1_)^1/2^). On this line, the growth is pure exponential with *α* = *μ*. But even if the system has reached this line before division, partitioning stochasticity will throw it off this line after division, because the *X* and *Y* populations of the daughter cell are chosen independently of each other at partition and will no longer have the same ratio as before. Thus in the next generation the trajectory will again be a superposition of two exponentials, resulting in a value of *α* = *μ*.

**FIG. 8:**
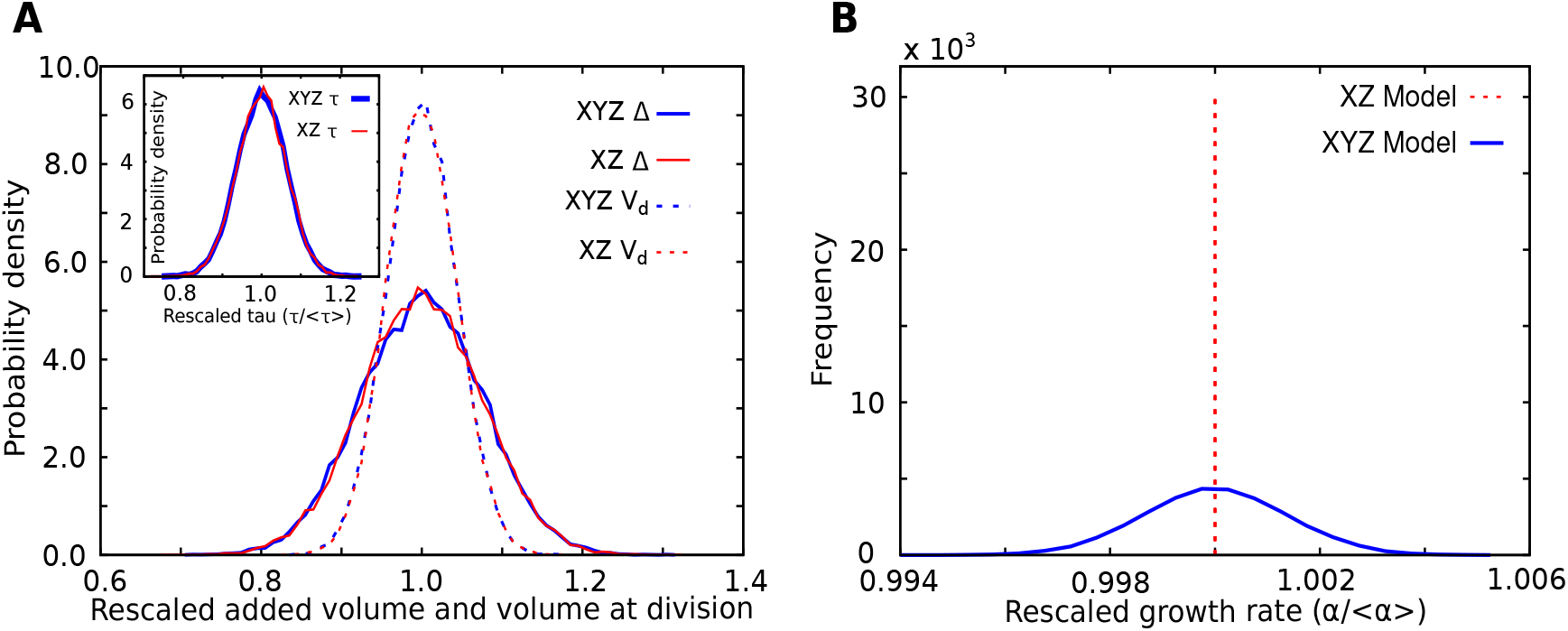
Distributions for XZ and XYZ models with only partitioning stochasticity. **A** Rescaled Δ, V_d_ distributions (*τ* distributions in the inset). **B** Rescaled *α* distribution. Distributions are made from 30000 generations. Note that the *α* distribution has zero width for the XZ model but nonzero width for the XYZ model, which explains how being thrown off the CBG contributes to the width of the *α* distribution (see discussion in main text). In these simulations intrinsic stochasticity due to chemical dynamics as well as threshold stochasticity are both absent, but partitioning stochasticity is present with strength *s* = 1. XZ model parameters: μ = 10, *K_Z_* = 10^-3^, *v* = 1, *s* = 1. XYZ model parameters: *c*_1_ = 10, *c*_2_ = 1, *K_Z_* = 10^-3^, *v*_1_ = *v*_2_ = 1, *s* = 1. For both models *Z_c_* = 40, *Z_r_* = 20, *τ*_1_ = 0.

The above argument makes it clear how being thrown off the CBG contributes to the width of the *α* distribution, because in every generation the departure of *X*(0), *Y*(0) (birth coordinates of the cell) from the CBG will be random due to partitioning stochasticity and the extent of that departure will govern how different the fitted value of *α* in that generation is from *μ*. Whereas in the XZ model since there can be no departure from the CBG partitioning stochasticity always gives *α* = *μ*. This explains why in the PTRZ model we found the width of the *α* distribution to grow with *s*: larger *s* means a greater departure from the CBG and hence a greater departure from a pure exponential solution. This also explains why when only threshold stochasticity is present (intrinsic stochasticity and partitioning stochasticity absent) the width of the *α* distribution is zero. This is because once the system reaches the CBG attractor, threshold stochasticity by itself cannot throw it off the CBG because it only operates in the *Z* sector and does not affect the ratios of populations in the *X* sector. In view of the above discussion it is not surprising that the *α* distribution depends upon parameter values and details of the models.

The mixing of exponentials as a possible origin of the *α* distribution has also been discussed in [61] in the context of a linear model. Our explanation, in terms of the cell being thrown off the CBG by partitioning and intrinsic stochasticity covers both linear and non-linear dynamics.

The above discussion also clarifies the origin of the departure from the adder property of an auto-regulated chemical *Q* discussed in the previous subsection. Since the cell is thrown off the CBG, the growth rate of *Q* is no longer *μ* but can be affected by *Q_b_* because of autoregulation. Hence it departs from the adder. PTR also have a growth rate different from *μ*, but being large and unaffected by *Q*, do not show a significant departure from the adder property (and hence V also does not).

## V. THE ORIGIN AND CONSEQUENCES OF EXPONENTIAL GROWTH

### A. Exponential growth arises in Class-I dynamical systems

We saw that the PTR sector defined by (2) and (3) was characterized by the exponential growth of chemicals in the deterministic steady state in the growth phase (insets of Figs. 1A,C; Fig. S9A,B), given by Eq. (7). We now argue that this is a general property of a large class of chemical systems which naturally arise in cellular dynamics. Consider a set of n chemicals whose dynamics is given by (1a), with *f_i_* being *homogeneous degree-1* functions of the molecular populations, i.e., for any *β* > 0,

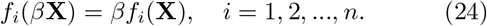

We will refer to systems satisfying (24) as *Class-I* dynamical systems. For such systems an exponential ansatz

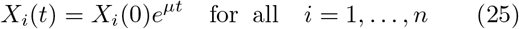

is generically a solution of the dynamics. Substituting equation (25) in equation (1a), and using equation (24) to write

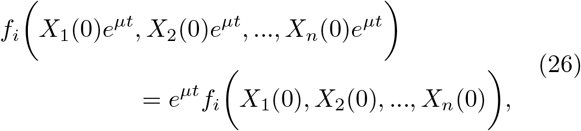

we get:

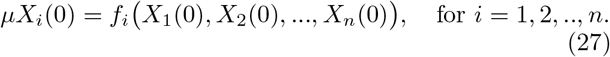

The *t* dependence has canceled out from both sides because of the class-I nature of the *f_i_*. This shows that for class-I systems (25) is a solution of the dynamics if and only if **X**(0) and *μ* satisfy (27). (27) is a set of *n* equations for the *n* + 1 constants *μ* and *X_i_*(0) appearing in (25). Assuming *X_n_*(*t*) > 0, we can define the ratios of populations *ψ_i_*(*t*) ≡ *X_i_*(*t*)/*X_n_*(*t*), *i* = 1, 2,…, *n* − 1, *ψ_n_* ≡ 1. It follows that *ψ_i_* satisfy the differential equation *dψ_i_*/*dt* = *f_i_*(*ψ*) − *ψ_i_f_n_*(*ψ*), where *ψ* denotes the vector (*ψ*_1_, *ψ*_2_,…, *ψ*_*n*−1_, 1). Under the ansatz (25) *ψ_i_* are time independent, *ψ_i_*(*t*) = *ψ_i_*(0) = *X_i_*(0)/*X_n_*(0). Dividing both sides of (27) by *X_n_*(0) and using (24), we get

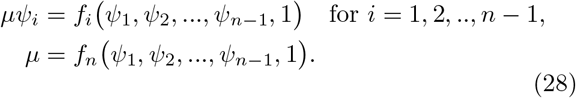

For class-I systems (27) and (28) are equivalent. (28) is a set of n (in general, non-linear) equations for the n unknowns *μ* and *ψ*_1_,…, *ψ*_*n*−1_. Generically these n equations provide a solution for the *n* − 1 independent ratios of chemicals *ψ_i_* (*i* = 1,…, *n* − 1) as well as *μ* in terms of the parameters appearing in the functions *f_i_* (see Supplementary section S2 D for additional remarks and qualifications). The PTR model is a particular case of *n* = 3 for which the explicit solution is given in (8). The fixed values of the ratios *ψ_i_* so obtained define a straight line passing through the origin of the n-dimensional phase space Γ of the variables **X** in the direction of the vector *ψ*. The implication of the above analysis is that for any initial condition **X**(0) lying on this line, the trajectory of system will satisfy (25); in other words all the populations *X_i_*(*t*) will grow exponentially with the same rate *μ* preserving their ratios. Therefore this line can be referred to as a *curve of balanced growth* (CBG) for the system (a trajectory that starts on the CBG remains on the CBG with constant ratios of populations). Such a curve does not exist in general for systems that are not class-I.

As discussed earlier in sections IV A and IV B and Fig. S9, in the case of the PTR dynamics defined by (2) and (3), an exponentially growing trajectory lying on the CBG is not just a solution but also a stable attractor of the dynamics. We have found this property to be true in simulations of many other class-I chemical systems representing cellular dynamics with different forms of the functions *f_i_*. This includes autocatalytic systems with regulation involving Hill functions (see Eq. (22) and models in [62]), another with a network of a thousand chemical species constructed along the lines of [63], and other complicated class-I autocatalytic systems (details to be presented elsewhere). On the other hand, without the class-I property, even simple systems do not have an attractor satisfying (25). For example we can consider the PTR model with the same Eq. (2) but with a modified Eq. (3) such that the volume is not a linear function of the populations. Let us assume that *V* is proportional to *S*^2/3^ where *S* is the surface area of the cell (motivated by spherical shaped cells), and let us take *S* to be proportional to *T*, the number of transporter molecules. Then the asymptotic trajectory is not exponential (see Fig. S10). The reintroduction of the division process does not produce the exponential behaviour in this non-class-I system. Similar behaviour is observed in several other non-class-I systems we have studied.

Thus we find the class-I condition (24) to be an important condition for the existence of attractors with exponential growth. Note that linear functions (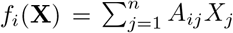 with constant *A_ij_*) are a special case of class-I and here the asymptotic attractor is exponential: **X**(*t*) = **X**^(*μ*)^*e^μt^* where *μ* is the eigenvalue of the matrix *A* = (*A_ij_*) with the largest real part and **X**^(*μ*)^ the corresponding eigenvector. In this case *ψ* is proportional to **X**^(*μ*)^. Class-I systems are in general nonlinear (e.g., the PTR model above) but share the feature of exponential solutions with linear systems. (27) can be considered a generalization of the eigenvalue equation *A***X** = *μ***X** to the nonlinear case, which determines the ratios of the *X_i_*(0) and the ‘eigenvalue’ *μ*.

### B. When does the class-I property arise?

The class-I property of *f_i_* will always arise for well-stirred mass action kinetics, however complicated the latter may be, when the volume of the container is a linear function of the chemical populations. This can be seen as follows.

Let us consider a well stirred chemical reactor of fixed volume *V* containing *n* chemical species whose populations are given by *X_i_* and concentrations by *x_i_* = *X_i_/V*. The law of mass action implies that the dynamics of the concentrations is given by the set of nonlinear equations

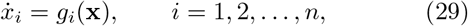

where *g_i_* are some nonlinear functions of the arguments that depend upon the set of chemical reactions that take place in the system. E.g., if there is a chemical reaction of the kind A + 2B → C, then *ẋ_C_* will contain a term of the kind 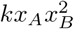. It is important to note that the r.h.s. of (29) is a function of the concentrations of the chemicals, not their populations.

Now assume the container is expanding, with the volume having a time dependence *V* = *V*(*t*). Then the above equation would be modified to

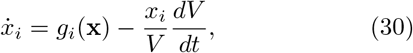

with the second term reflecting the effect of dilution due to expansion. Let us ask for the dynamical equations in terms of the populations *X_i_* = *Vx_i_*. In a fixed (constant) volume, *Ẋ_i_* = *Vẋ_i_* = *Vg_i_*(*x*) = *Vg_i_*(**X**/*V*). In an expanding volume 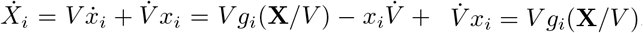. Thus in terms of the population variables the dynamics in the expanding container does not contain any extra term and is given by

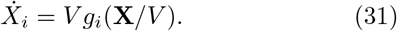

Now suppose the volume of the container is a function of the populations, *V* = *V*(**X**), i.e., it depends upon time only through **X**(*t*). This feature would modify the nature of the dependence of the r.h.s. of (31) on **X**:

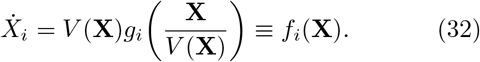

A particularly interesting situation arises when *V*(**X**) is a linear function of the populations (as is possibly true for bacterial cells):

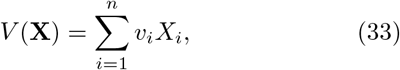

where *v_i_* are constants. (This can happen for example due to osmotic pressure. If we assume that water enters or leaves the cell on a short time scale compared to the time scales of the dynamics (31) to maintain the total concentration of solute inside equal to that outside the cell *x_ext_*, then 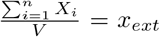 or 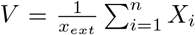. This is a particular case of (33) where the sum over *i* includes all the chemical species in the bulk (interior) of the cell.) Then it follows that *f_i_* satisfies the class-I property. For, (33) implies that *V*(*β***X**) = *βV*(**X**); then 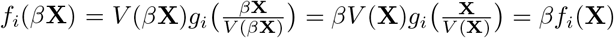.

Thus the class-I property (24) follows from well stirred mass action kinetics in an expanding container Eq. (30), and the assumption that *V* is itself a homogeneous degree-1 function of the molecular populations (in particular its special case (33) that it is a linear function of the populations). Note that the *g_i_*(**x**) can be highly nonlinear functions of their arguments, and so will in general *f_i_*(**X**) be; nevertheless the *f_i_* will satisfy the condition (24). This property of well stirred mass action kinetics is hidden when the dynamics is formulated in terms of concentrations but is apparent when formulated in terms of populations which are extensive quantities.

Note that the dynamical system (31) in terms of extensive variables is not fully specified until *V* is specified as a function of time or as a function of **X**. Similarly the dynamical system (30) in terms of intensive variables is not fully specified until 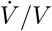 is specified as a function of time or as a function of **x**. The choice (33) specifies both dynamical systems completely where the constants *v_i_* are treated as parameters of the system. With this choice the concentrations satisfy the constraint 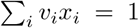, obtained by dividing both sides of (33) by *V*. It can be easily seen that this constraint is preserved by the time evolution under (30). Thus there are only *n* − 1 independent intensive variables, which can be taken to be the *n* − 1 independent *x_i_* or the *n* − 1 *ψ_i_* (each set can be expressed in terms of the other set).

It is important to note that the specification (33) allows us to find the steady state growth rate as a function of the parameters. (33) implies that the *f_i_* appearing in (32) satisfy the class-I property and this leads to (27) under the ansatz (25) from which both *μ* and the *ψ_i_* (or *x_i_*) can be determined. The x so obtained is a fixed point of (30). The *μ* and x so obtained depend, among other things, on the parameters *v_i_* appearing in (33) (see, e.g., (8c)). If, instead of specifying *V* as a function of **X**, we had simply replaced 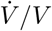 by *μ* in (30), we could still solve for the fixed point of (30) in terms of *μ* and other parameters appearing in the functions *g_i_* but we would not be able to solve for *μ* in terms of the parameters.

### C. Consequences of the class-I property

The class-I property allows us to generalize the results of the PTRZ model to a much more general class of models described in section II. In this subsection we discuss the generalizations and the assumptions under which they hold.

#### 1. Exponential growth of size, expressions for the average birth volume and interdivision time scale

##### Exponential trajectories

Most mathematical models of cellular dynamics are formulated in terms of concentrations, with particular choices of the functions *g_i_* in (29). For cells in a steady state culture it is usually *assumed* that the volume grows exponentially; consequently a term − *μx_i_* is added to account for dilution due to volume expansion. However, the exponential dependence of *V* on *t* is, *a priori*, a puzzling fact given the nonlinear nature of cellular dynamics. The discussion above provides an explanation of that, and also explains why the exponential growth property is so generic and independent of the form of *g_i_*. As remarked in section *V* A we find exponentially growing chemical trajectories (25) as attractors of the dynamics for a wide range of systems irrespective of the form of *f_i_* when the *f_i_* are class-I. In all these cases *f_i_* were derived from physically motivated g¿ via Eqs. (32) and (33). Then, since the volume of the container is a linear function of the populations (33), it follows that the container size in the attractor also grows exponentially with the same rate: *V*(*t*) ~ *e^μt^*. Thus the ubiquity of exponential growth of *V* is a consequence of the fact the *f_i_* are class-I irrespective of the form of *g_i_*, and that such systems possess exponential solutions which are often the attractors of the dynamics.

This explanation also exposes a physical property that is required for exponential trajectories that has not been so far recognized, namely the linear dependence (33) of *V* on the populations (or, more generally, *V*(*β***X**) = *βV*(**X**)). This suggests that the biophysical origins of the assumption (33) need to be explored further. In particular the constants *v_i_*, which also affect the steady state growth rate of the cell (and hence cellular fitness), should be determined experimentally and estimated theoretically. This also provides a possible explanation of the departure from exponential trajectories observed in certain eukaryotic unicellular organisms [4, 13]. The departure may be in part a consequence of the violation of (33), caused by other structural features of the cell such as the cytoskeleton.

##### Expression for *τ*

Let us consider the consequences of this for the general model described in section II. We assume, for the rest of this section and the next section that **(a)** *V* is a linear function of the populations and the functions *f_i_* satisfy the class-I property, and **(b)** that an exponential solution (25) is the attractor of the *X* sector dynamics for the initial conditions of interest. In discussing averages we ignore all forms of stochasticity and treat the dynamics as deterministic. Then, the *X_i_* variables flow towards the CBG defined by the vector *ψ* whose components, the ratios of populations, as well as *μ*, satisfy (27). This flow is not affected by the change of *Z* because *Ẋ_i_* is independent of *Z*, except at discrete points when all the *X_i_* are halved. The latter interruption does not disturb the flow towards the CBG because 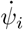 are unaffected by it (as mentioned above (1a) and (24) imply that 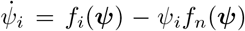, which is invariant under change of scale of the *X*). The dynamics of ratios of chemicals and their concentrations (30) therefore does not see any discontinuity at division. Thus the *X* variables reach the CBG in the presence of the division process. This will be true as long as division changes all the *X_i_* by the same scale (not necessarily half), and irrespective of whether division is triggered by the *Z* dynamics as in the present case, or by some other process. Thus, in particular it is guaranteed that in the attractor of the growth-division dynamics, the cell volume and all populations *X_i_* will grow exponentially with time between birth and division. Furthermore in the steady state, since the variables must grow by a factor of 2 between birth and division (assuming *X_i_* are halved at division), the interdivision time is given by *τ* = ln 2/*μ*. Note that *τ* only depends upon the parameters of the *X* sector and not the *Z* sector.

##### Expression for *V_b_*

Now let us further assume that **(c)** the function *h*(**X, Z**) in (1b) is independent of *Z* and satisfies the class-I property, *h*(*β***X**) = *βh*(**X**) (as is the case in the PTRZ model, Eq. (5)). When the *X* have reached their CBG we can write 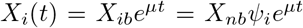 or 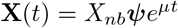, where *X_ib_* is the value of *X_i_* at a birth of the cell (taken to be *t* = 0 for convenience). Then *ṻ* = *h*(**X**) = *Ce^μt^*, with the last step following from (c), and *C* = *X_nb_h*(*ψ*). This is the same as (10), except that the expression for *C* is now more general. The argument below (10) and in the Supplementary text section S2 A applies as before, fixing the absolute scale of the *X_ib_* and *V_b_*. In particular *X_nb_* = *Nμe*^*μτ*_1_^/*h*(*ψ*) and since 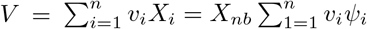, we get

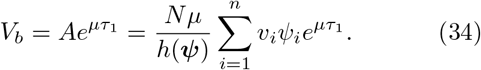

This is the expression for cell size at birth in terms of the parameters for the general model described in section II when the conditions (a)-(c) hold. (13), (14) for the PTRZ model are special cases of this formula. Note that *μ* and *ψ_i_* are determined in terms of the parameters appearing in the *f_i_* through (27).

#### 2. Balanced growth of chemicals

In order to replicate itself a growing cell must solve a high dimensional coordination problem between its chemicals. The amount of each chemical in the mother cell at division must be double that in the daughter cell at birth (assuming that upon division a daughter gets half of every chemical from its mother). Otherwise daughters at birth in successive generations will not be identical. How does the cell manage to double all of its chemicals (and there are thousands of them) at the same time? This is a puzzle because each chemical is produced and consumed in a specific set of reactions that have their own specific rate constants, varying from reaction to reaction. However if the exponential solution (25) is an attractor of the dynamics (1a) the problem is automatically solved because in the attractor each chemical has the same growth rate μ and ratios of chemicals are preserved in time. It is remarkable that class-I systems describing cells seem to have such attractors. This is not true for systems that are not Class-I (for an example, see Fig. S10), which must solve their coordination problems by other means (to be discussed elsewhere).

#### 3. Genericity of the adder property and the Δ and t distributions

So far in this subsection we have discussed the consequences of the class-I property for the general system (1) at the deterministic level (without stochasticity). Now consider the inclusion of intrinsic stochasticity in the chemical dynamics of *Z*, while still treating the dynamics of the X sector chemicals as deterministic, and also ignoring all other sources of stochasticity such as partitioning stochasticity, threshold stochasticity, etc. The adder property and the Δ and *τ* distributions discussed in section IV D derived from three key ingredients: (i) *V*(*t*) = *V_b_*e^*μt*^, (ii) cell division occurs at the time when *Z* first reaches *Z_c_*, whereupon it is reset to a fixed value, and (iii) the dynamics of *Z* is the stochastic version of (10) in which *C* is an extensive quantity proportional to the chemical populations and hence to the volume of the cell. In the class of models described in section II property (ii) is taken for granted (it is part of the definition). Property (i) follows from assumptions (a) and (b) mentioned above (in section VC1). As noted above, when assumption (c) holds, then on the CBG attractor of the *X* sector deterministic dynamics, (10) also holds with *C* and *V_b_* both proportional to *X_nb_* and hence to each other. Thus (iii) also holds provided we treat the *Z* dynamics as stochastic. This proves that under conditions (a)-(c) when we treat the *X* sector dynamics as deterministic and the *Z* dynamics as stochastic the general model described in section II displays the adder property and the Δ and *τ* distributions given in section IV D (Eqs. (15) to (18)) arise with *C* = *h*(X_*b*_) and *B* = *C/μ*. Further, the populations X_*b*_ (rescaled by their means) also show the same distribution as *V_b_* and exhibit the adder property.

When the *X* sector dynamics are also treated stochastically, and other sources of stochasticity such as partitioning stochasticity and stochasticity of *Z_c_* are included, we expect that the model will exhibit a behaviour similar to that discussed for the PTRZ model in section III B, e.g., a non-trivial *α* distribution will arise, the adder property for the cell volume will continue to hold unless Zr and Zc are correlated, the CV of molecular populations as a function of the mean 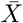 will decline as 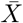 increases for small 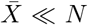 and flatten out for 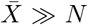, etc.

## VI. UNIFIED DESCRIPTION OF BACTERIAL CELL SIZE BEHAVIOUR AND GROWTH LAWS

So far we have discussed issues related to the size of bacterial cells in a growth culture. The models considered here have a modular character, in that there are two sectors, the *Z* sector which is concerned with the triggering of division, and the *X* sector which contains all other chemicals in the cell. We have seen that certain properties of the models are independent of the details of the *X* sector, as long as certain broad conditions are met (class-I nature of dynamics, linear dependence of *V* on populations, etc.). The average cell volume at birth depends upon various cellular parameters (13),(14),(34). It depends on the *X* sector parameters through the growth rate μ, the ratios of chemicals *ψ_i_* and the coefficients *v_i_*, and on the *Z* sector parameter *N/K_Z_* (or *N/h*(*ψ*) in general). It also has the exponential factor *e*^*μτ*_1_^. The overall dependence of *V* on *μ* is complicated in light of the fact that the ratios *ψ_i_* also depend upon the same parameters of the *X* sector that *μ* depends upon (through a solution of (27)) and this would be different for different models. On the other hand the fluctuations in *V* and the populations *X_i_*, in particular their CV, as well as the CV of *τ* are governed by a single parameter *N*, and are largely independent of the details of what happens in the *X* sector. Experiments seem to constrain the parameter *N* between 20 and 60, as discussed in section III B 4. The properties of exponential growth and the adder property for *V* and *X_i_* are also largely independent of the details of the *X* sector.

We now mention another aspect of bacterial growth physiology that pertains to the effect of the medium on the growth rate and composition of the cells, the bacterial growth laws. These are summarized in three empirically derived equations:

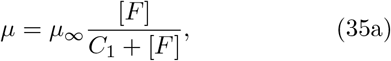

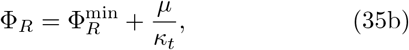

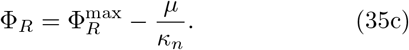

The first [64] describes the dependence of the steady state growth rate on the concentration [*F*] of a limiting food resource in the external medium. The second [40–43] describes how the fraction of total protein in the cell that is ribosomal protein, Φ_*R*_, increases as *μ* is increased by improving the nutritional quality of the limiting food resource. The third [43] describes how Φ_R_ increases as μ is *decreased* by adding antibiotics to the medium that diminish the translational efficiency of ribosomes or by producing mutants that specifically target the translational efficiency. The six constants 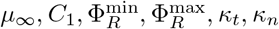 are phenomenological constants [43] whose values are extracted from experiment.

In [50] the above three laws were derived from the PTR model, which is the *X* sector of the PTRZ model discussed above. There are two classes of macromolecules represented in the PTR model, *T* and *R*, made up of *m_T_* and *m_R_* units of *P* respectively, and the ribosomal fraction can be defined as Φ_*R*_ = *m_R_R*/(*m_T_T* + *m_R_R*) = *m_R_*/(*m_T_* + *m_R_*). To derive the growth laws (35) the expression (8c) for the steady state growth rate was maximized with respect to *f_R_*, keeping all other parameters fixed. This implements an underlying regulatory mechanism in the cell that regulates the fraction of ribosomes that are engaged in making ribosomal protein. The value of f_R_ that maximizes the r.h.s. of (8c) was substituted in (8a) to get an expression for (and hence Φ_*R*_) and in (8c) to get an expression for μ. This yielded the equations (35) for the growth laws together with expressions for the six phenomenological constants in terms of the parameters of the PTR sector. For the parameter values given in Fig. 1 the predictions of the model agree with the experimental data up to factors of order unity.

It is clear that the above procedure and its results are unaffected by the presence of the *Z* sector in the PTRZ model. The modular structure of the PTRZ model and the fact that the division process does not cause any discontinuity in the dynamics of the intensive variables of the *X* sector ensures that the PTRZ model also reproduces the same equations for the bacterial growth laws. Setting *f_R_* to a particular value (that optimizes *μ*) affects the values of *μ* and the ratios and *ψ_R_*, and hence the value of Vb in (14), but does not affect the existence of exponential growth. In particular (10) still holds albeit with values of *C* and *μ* given by the above procedure. Thus the consequences of the *Z* sector dynamics are also unaffected. The PTRZ model thus provides a unifed explanation of the bacterial growth laws together with fluctuations of size, interdivision time, growth rate and intracellular molecular populations as well as the adder property.

The PTR sector can be augmented [62] by introducing other chemical species whose dynamics model the regulation of *f_R_* mechanistically instead of using the optimization procedure. Standard regulatory mechanisms are consistent with the general form (29) and do not alter the class-I nature of the *X* sector. The feature of exponentially growing trajectories remains intact in such models. The results above about cell size and fluctuations will therefore also hold for such models.

## VII. SUMMARY AND DISCUSSION

We have presented a class of mathematical models that explain a number of observed properties of bacterial cells and make testable predictions. The work also introduces new concepts that may help further theoretical analysis and identify new experiments.

### Summary of assumptions and results

The models assume that the cell can be described in terms of the intracellular chemical populations *X_i_, i* = 1,…, *n*, whose growth dynamics, at the deterministic level, is given by coupled ordinary differential equations (1a). The cell volume *V* is assumed to be a linear function of the chemical populations, Eq. (33). The models have further structure to describe the control of cell division, but already, even without that structure, certain key properties of the cell, summarized in the next paragraph, are fixed by the ‘X sector’ itself. These are governed by the character of the functions *f_i_* appearing in (1a).

#### Class-I property: Exponential trajectories, growth rate and interdivision time scale, intensive quantities

The models provide an explanation of the observation that cell size and intracellular molecular populations grow exponentially with time between birth and division in steady state bacterial cultures. Exponential growth is shown (in section V A) to be a consequence of the ‘class-I’ property (Eq. (24)) of the functions *f_i_*. This property in turn is a general consequence of mass action kinetics in expanding containers (see section *V* B) when the cell volume *V* depends linearly on the populations (Eq. (33)) and could therefore hold for a large class of models applicable to bacteria. When the class-I property holds, we find that the populations are typically attracted to a ‘curve of balanced growth’ (CBG) in which their ratios *ψ_i_* ≡ *X_i_*/*X*_*n*_ are constant, and the populations themselves grow exponentially in time (Eq. (25)) with a constant growth rate μ. This explains the phenomenon of balanced growth [18] in bacterial cells. This also explains why *V* grows exponentially in the steady state. The ratios *ψ_i_* and *μ* are determined in terms of the constant parameters of the model as solutions to a set of (typically algebraic) equations (Eqs. (27)), which constitute a nonlinear analogue of the eigenvalue equation of a matrix. Hence all the intensive quantities in the steady state including the concentrations *x_i_ ≡ X_i_/V* of the chemicals are fixed. The interdivision time scale *τ* is fixed by the reciprocal of *μ*. E.g., if the division is symmetric (i.e., the two daughters are identical), then *e^μτ^* = 2 since all chemicals should double in quantity between birth and division in the steady state; hence *τ* = ln 2/*μ*. Explicit solutions for *μ* and *ψ_i_* (Eq. (8)) are presented for a nonlinear model with *n* = 3 populations, the PTR model [50] defined by Eqs. (2). We also show examples of non-class-I models where the growth trajectory of the individual cell is not exponential (section VA, Fig. S10).

#### Bacterial growth laws

The PTR model also reproduces the bacterial growth laws of composition [40–43] when regulation is implemented through an optimization procedure [50] or by introducing additional molecular species [62] while preserving its class-I property and exponential trajectories (section VI).

#### Division control: Absolute size and populations

While intensive quantities and the interdivision time scale in the steady state are fixed by the previous assumptions, the absolute scale of cell size and populations requires a specification of the division control mechanism. In this work we assume that the cell commits itself to division when one of the chemical populations (denoted by *Z* and separated out from the *X_i_* for convenience) reaches a threshold *Z_c_*, with division following commitment after a possible delay *τ*_1_. The dynamics of *Z* depends upon the *X_i_* through (1b). For simplicity we assume that *Z* contributes negligibly to its own dynamics, to the continuous time dynamics of the *X_i_* and to *V*. Immediately after it reaches its threshold *Z_c_* we allow *Z* to be reset to a value *Z_r_* < Z_c_. Upon division all the populations are halved. With these assumptions division is governed by the nature of the function h appearing in (1b). In this paper we have primarily considered the consequences of *h* also satisfying the class-I property like the *f_i_* (Eq. (24)). Then we can get a general formula for the birth volume Vb (Eq. (34)) in terms of h and a specific formula in the case of the PTRZ model (Eq. (14)) when a specific form (given by (5)) of h is chosen. The absolute populations are also fixed (expressions given above (34) and (13)).

#### Stochasticity in chemical dynamics of Z: Analytic distributions of τ and Δ; the adder property for cell size and chemical populations

The preceding two paragraphs summarize results for deterministic versions of the models, and therefore pertain to averages across cells in the cultures. In order to understand cell-to-cell variation in steady state cultures we include various sources of stochasticity. Our stochastic results are obtained for the case *τ*_1_ = 0. It is convenient to distinguish four sources of stochasticity that we have considered (which have distinct physical consequences): (A) Intrinsic stochasticity in the chemical dynamics of *Z*, (B) intrinsic stochasticity in the chemical dynamics of the *X* sector chemicals, (C) partitioning stochasticity, and (D) stochasticity in the value of the threshold *Z_c_*. Intrinsic stochasticity results from the fact that each chemical reaction is a molecular event with a certain probability. In particular this makes the inter-division time, which by assumption is the first passage time for *Z* to reach *Z_c_*, a stochastic quantity. Our implementation of (C) and (D) is defined in section IIIB 5. We have investigated different types of stochasticity in isolation and in suitable combinations. When type A is the only stochasticity present we obtain some analytic results for general class-I systems, namely, the distributions of *τ* (15), and of added volume Δ (16), both conditional on the birth configuration of the cell. The conditional Δ distribution is independent of *V_b_*, thereby proving the adder property. In fact the rescaled Δ, *u* ≡ Δ/〈Δ〉, has a distribution (18) that depends on only one parameter, *N = Z_c_* – *Z_r_*/2 and is independent of the details of the functions *f_i_, h* and other parameters (section VC3; Fig. S1). Molecular populations also satisfy the adder property and their rescaled increments have the same distribution as *u*. However, the distribution of growth rate *α* has zero width when only type A stochasticity is present (discussed in the beginning of section IV D).

#### Other sources of stochasticity: Robustness of size and τ distributions; cross-over of population distributions; origin of the growth rate distribution; departure from the adder property

We have studied stochasticities of type B,C,D with numerical simulations of the nonlinear PTRZ model (defined by Eqs. (2)(3)(5)) and two linear models, the XZ and XYZ models (Eq. (23)). We find, in the parameter ranges considered, that the rescaled distributions of Δ, *τ, V_d_, V_b_* obtained with purely A type stochasticity are robust to the inclusion of B type stochasticity (Figs. S17), and also to the inclusion of the C and D type stochasticities (section IIIB 5; Figs. S5) provided their strength is not too large. The robustness of the rescaled size and interdivision time distributions to parameter values (except *N*) and other sources of stochasticity is one of the striking results of this work. The CV of all these distributions is proportional 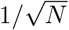 when only the A and B type stochasticities are included (Fig. 2C), with proportionality constants given by Eq. (6). The rescaled distributions of the populations *X_i_* are also robust, provided the population mean 〈*X_i_*〉 is not too small. We find a crossover behaviour of the CV of *X_i_* as a function of 〈*X_i_*〉 (Fig. 5), with CV ~ 〈*X_i_*〉^−1/2^ for 〈*X_i_*〉 ≪ *N* and CV constant for 〈*X_i_*〉 ≫ *N*. The adder property of the volume and populations is also found to be robust to the introduction of the BCD type stochasticities (Figs.4, S4, S6A). However the adder property is lost with D type stochasticity if *Z_c_* and *Z_r_* are correlated with each other (Fig. S6B). The distribution of growth rate *α* acquires a non-zero width upon the introduction of type B stochasticity (Fig. 2), depends upon parameters other than *N* (Fig. S1) and is strongly influenced by type C stochasticity (Fig. 6). We show that the origin of the *α* distribution is in part a consequence of the fact that type B and C stochasticities throw the populations off the CBG and thereby cause a mixing of a pure exponential function of time (having a rate *μ*) with other functions of time (including exponentials with rates other than *μ*).

#### Comparison with experiment

At the level of averages, the models reproduce the exponential trajectory of the volume and intracellular populations as seen in *E. coli* experiments, as well as the bacterial growth laws of composition. At the level of fluctuations, they reproduce the adder property of cell volume fluctuations in *E. coli*. Experimental distributions of Δ, *τ, V_d_* [11] constrain the parameter *N* of the models to be between 20 and 60 (section IIIB 4). The models find a much narrower distribution of *α* than the one observed in [11] when only the intrinsic stochasticity of chemical dynamics is included, and require a significant strength (perhaps unnaturally large) of partitioning stochasticity to reach the observed width (section IIIB 5). This discrepancy needs to be explored further, both theoretically and experimentally. The models reproduce the observed crossover behaviour of the CV of a molecular population as a function of its mean population 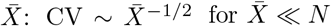 and constant for 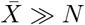 (section IIIB 3).

### Discussion

A key question is: Which molecular population in the cell does *Z* correspond to? One possible candidate is the DnaA molecule, which is known to initiate DNA replication and has been suggested as an upstream trigger for cell division (for reviews see [65–68]) DNA replication in *E. coli* is known to be initiated when a certain number, believed to be between 20 and 30, of active (ATP-bound) molecules of DnaA bind to sites on the DNA molecule at *oriC*, a region of DNA at the origin of replication. Soon after the initiation of replication the DnaA is deactivated by other enzymes; the complex of active DnaA falls apart to prevent multiple rounds of initiation. Each initiation is followed first by the replication of DNA (referred to as the C period in the bacterial growth cycle) and then by the separation of the chromosomes into two halves of the cell and cell division (referred to as the D period). After division, the daughter cells have a smaller number of active DnaA molecules bound to the above mentioned sites, and this number grows in the period between birth and the initiation of replication (known as the B period). Various aspects of the dynamics of DnaA have been modeled mathematically [67, 69–72].

In the context of the present model it is tempting to identify *Z* with the number of active DnaA molecules in the initiation complex bound to *oriC*. Then present experiments with DnaA suggest that the parameter *Z_c_* of the model should be between 20 and 30, *Z_r_* should be essentially zero (the initiation complex dissociates after triggering replication) and *τ*_1_ = *C* + *D* ~ 1 hr. It is interesting that this identification leads to a value of *N* between 20 and 30, which overlaps with the range 20-60 obtained from a completely independent experimental constraint, the spread in the distributions of Δ, *τ, V_d_*. In other words, this identification provides a natural explanation of why the CVs of cell size, interdivision time and the large intracellular molecular populations are in the ballpark of 20% (Eq. (6) with *N* ~ 25). The model predicts that the average cell size increases when the rate of production of the *Z* population is lowered (*V_b_* is inversely proportional to *K_Z_* or *h*(*ψ*), Eqs. (13),(34)). This is consistent with the empirical observation that cell size increases when the production of DnaA is impaired [73]. We note that the fact that cooperativity in the *Z* dynamics leaves the adder property of the volume intact (shown in section IV D 2 and Supplementary section S2 C) is an encouraging sign for the above interpretation of Z, as such cooperativity is known to exist for the active-DnaA molecules bound at *oriC*.

However, the model also has problems with respect to the above interpretation. Note that in the previous para, *Z* is identified not with the total number of DnaA protein molecules in the cell, or even the total number of active (ATP-bound) DnaA molecules in the cell, but with the number of active DnaA molecules *bound to the DNA molecule at oriC.* A problem with the present model is that a production term for *Z* such as in Eqn. (5) where 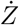 is an extensive quantity may be appropriate for a chemical produced in the bulk of the cell, but for a chemical species localized in space (such as the set of active-DnaA molecules bound to the DNA at *oriC*) would need further justification or a mechanism not provided by the present model. Further, in this interpretation it is not clear how to account for the synchrony observed in the initiation of replication at multiple origins [74]. The resolution of the above mentioned problems might lie in the fact that *oriC* and the *dnaA* gene are close by on the DNA molecule and spatial proximity effects need to be taken into account. It also needs to be noted that the dynamics of active-DnaA molecules bound to the origin of DNA replication is affected by many factors in the cell, including autoregulation of DnaA production, binding of DnaA to a large number of sites other than *oriC* on the DNA and sequestering of the binding sites by other enzymes. Another proposed model [75] is that initially in the cell cycle DnaA binds other sites on the DNA and the replication initiating event is the binding of active DnaA to a site in *oriC* that triggers cooperative binding of active DnaA on a relatively short time scale to *oriC*. In this interpretation *Z_c_* would correspond to the effective number of sites that DnaA binds to before the cooperative binding event occurs. In the light of the above caveats, the question as to whether the *Z* population in the present model or its extension corresponds to some sub-population of the DnaA protein in the cell remains an open question to be investigated further.

Note that the adder property and the Δ distribution (16)(and consequently Eq. (6) relating the CV of Δ to 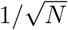) follow from very general assumptions (Eq. (19) and assumptions mentioned below it; or the class-I property and assumptions mentioned in V C 3). They and many other properties discussed above do not depend upon other details of the *Z* dynamics. Thus the models could apply to other candidates for the *Z* molecule than DnaA. Another candidate for *Z* is the FtsZ protein, which forms a ring around the cell that constricts and causes cell division, and degrades after cell division (for reviews see [76, 77]). However, the number of FtsZ molecules required for ring formation seems to be large (in the range ~ 10^3^ — 10^4^, suggesting that N is in the range ~ 10^3^ — 10^4^); in which case stochasticity in the production of FtsZ alone would not account for the observed CV of cell size and division time (the model predicts 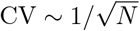 which would be too low). The present model ignores stochasticity in the mRNA population corresponding to the *Z* protein; its inclusion would enhance the CV of the cellular variables in (6) beyond 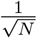 and in particular improve the case for FtsZ. Further, in the cell stochasticity in both DnaA and FtsZ can simultaneously contribute to the stochasticity in interdivision time and cell size.

In sum, while there are two good candidates (DnaA and FtsZ) for the time-keeper molecule, we believe that more work, both experimental and theoretical, is needed to establish whether or not the observed stochasticity in the interdivision time and cell size is a consequence of their populations (or appropriate sub-populations) having to reach a critical threshold. Of course, there may be other molecular candidates in the cell for triggering division that we are not aware of.

The models discussed in the present work predict that the adder property of the cell volume is accompanied by the adder property for coarse-grained intracellular molecular pools, such as the pool of amino acids in the cell or the pool of metabolic enzymes, whose copy numbers are ≫ *N*, and that the distribution of such a molecular population rescaled by its mean is the same as the rescaled volume distribution. Chemical population distributions have been measured and found to be universal [32] but the adder property for molecular pools has not been reported to our knowledge. Susman et al [61] have reported a departure from the adder property for two individual protein populations in single cell trajectories of *E. coli* but the adder behaviour of larger molecular pools (predicted by our models) remains an open empirical question. The models also predict that in the presence of threshold stochasticity (*Z_c_* being distributed over a range of values from cell to cell) the adder property is lost if the reset value *Z_r_* is correlated with *Z_c_*. It would be interesting if this could be tested by suitable mutations of *E. coli*, or by studying bacteria in which the adder property is absent.

An experimental question that this work draws attention to is the role of osmosis and other biophysical mechanisms in understanding exponential trajectories of cells. In section *V* B we argued that the linear dependence of Von the intracellular populations, Eq. (33), is a crucial requirement for exponential trajectories. As mentioned there such an assumption might be justified if cells equalize the osmotic pressure of solute inside and outside the cell. Cells actually maintain an osmotic pressure difference between their interior and the exterior. It is important to test whether the linearity assumption is valid and also to measure the coefficients vi. It is worth noting that these coefficients affect the steady state growth rate *μ* (see, e.g., Eq. (8c) and the expressions below it) and thus contribute to cellular fitness. A departure from the linear dependence of *V* on intracellular molecular populations, caused by other structural elements such as the cytoskeleton, may explain the departure from exponential trajectories observed in certain eukaryotic organisms.

At a mathematical level this work suggests that a certain class of dynamical systems, class-I systems (defined by (1a),(24)), are both generic and analytically useful for modeling bacterial cells. The present study shows that some physical properties (summarized above, and including averages and fluctuations) are universal for all such systems in that they do not depend upon the model and (many of) its parameters, and identifies some that are not. For this class the ‘steady states’ are exponentially growing trajectories whose growth rate *μ* is a solution of a nonlinear version of the eigenvalue equation, Eq. (27). When the functions *f_i_* are algebraic *μ* is given implicitly in terms of the model parameters as a solution to a set of algebraic equations. In this work we have only investigated some properties in a few examples of class-I systems; other mathematical properties and examples need to be investigated. Since exponential growth appears in many areas where the underlying dynamics is nonlinear, it is quite possible that these systems find applications elsewhere.

For any fixed environment *μ* is a measure of the organism’s fitness. Having it as a function of the system parameters allows us to describe the fitness landscape including the neutral directions, valleys and hills in parameter space. Thus it specifies the evolutionary paths in that environment. This class of models could be relevant for the study of evolution because they provide *μ* as a function of the parameters. It is worth mentioning that for the class of models studied here *μ* is independent of the parameters of the division control sector (e.g., the function *h* in (1b), and the thresholds *Z_c_* and *Z_r_*), which are therefore neutral directions of variation as far as steady state fitness is concerned. This is a consequence of a kind of modularity in the model implicit in the assumption that the dynamics of the *X* sector is independent of *Z* and is influenced by *Z* only through division. It would be interesting to empirically explore the extent of, or departure from, such a modularity in real cells.

Since the models discussed here are dynamical, they capture not just the steady state but also the transients. Thus they could also be useful in exploring those regulatory mechanisms that affect or seek to optimize performance over the transients. Extension of these models may find applications in modeling the stationary phase [78] and antibiotic environments [79] where the net cell population growth goes to zero with a balance between cell growth and death. It would also be interesting to use these models to make contact with other allometric modeling approaches [80] that seek to understand how the amounts of cellular components like ribosomes and proteins depend upon cell size across diverse bacteria.

## COMPUTATIONAL METHODS

All numerical simulations were done in *C* programming language. Numerical solution of the ODEs for the mathematical models were done using the CVODE solver library of the SUNDIALS (Suite of Nonlinear and Differential/Algebraic Equation Solvers) package [81] and the adaptive Runge-Kutta (RK5) method. The stochastic simulations were done using the tau leaping method [82, 83]. In particular the trapezoidal variant of adaptive implicit tau leaping was used [84, 85].

## Supporting information

## ACKNOWLEDGEMENTS

We would like to thank Mukund Thattai for a critical reading of an earlier version of the manuscript, discussions and helpful suggestions. We thank Suckjoon Jun for providing the data of ref. [11] for comparison of the model with experiment and for discussions. We also thank Santhust Kumar, Saurabh Mahajan, Amitabha Mukherjee and Pooja Sharma for discussions. SJ thanks the members of the Simons Center for Systems Biology, Institute for Advanced Study, Princeton for discussions and critical comments. He acknowledges the Addie and Harold Broitman Membership in Biology at the Institute for Advanced Study, grants from Department of Biotechnology, Government of India, and a Research and Development grant from the University of Delhi. PPP and HS would like to thank the University Grants Commission, India for a Senior Research Fellowship and Junior Research Fellowship, respectively. We would like to acknowledge the hospitality of the International Centre for Theoretical Sciences, Bangalore, and the International Centre for Theoretical Physics, Trieste, where part of this work was done.

