## Supplementary material for "Exponential trajectories, cell size fluctuations and the adder property in bacteria follow from simple chemical dynamics and division control"

### S1. SUPPLEMENTARY FIGURES

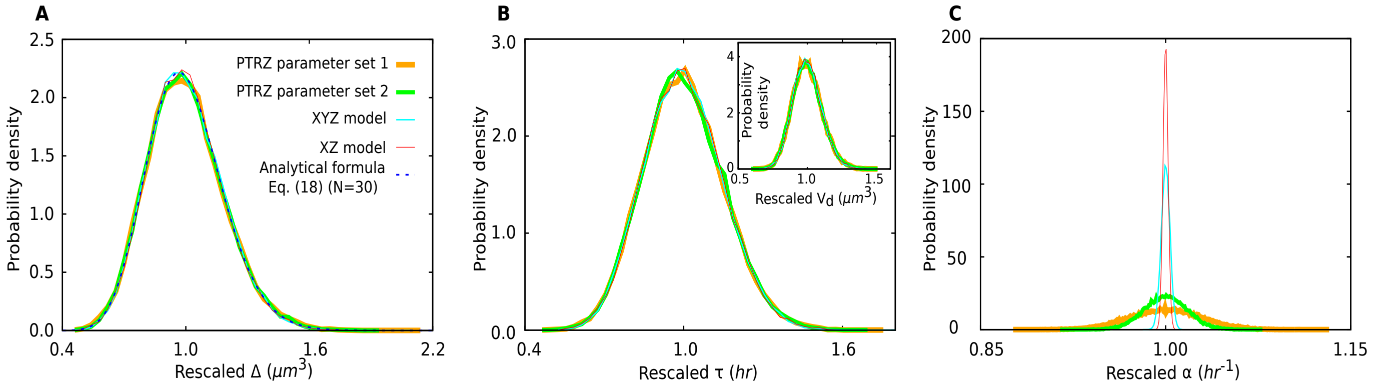


FIG. S1: **Robustness of the qualitative dynamical behaviour and rescaled distributions to parameter and model changes**. **A,B,C** Distributions of rescaled $\Delta$, $\tau$ and $\alpha$, with $V_d$ in the inset of **B** compared for two parameter sets of the PTRZ model and other models. The legend of the curves for different parameter sets and models in all three figures is given in **A**. We present results for a set of parameters of the PTRZ model that differ from those in Fig. 1 in growth rate, ratios of chemicals, averages of size, interdivision time, etc., and compare the results for the two sets. The parameter values of Fig. 1 are referred to as PTRZ parameter set 1 in the legend. The new parameter values (PTRZ parameter set 2) are $K_P = 5 * 10^5 hr^{-1}$, $k = 10^{-1} hr^{-1} (\mu m)^3$, $d_T = 0.01 hr^{-1}$, $d_R = 0$, $m_T = 4000$, $m_R = 100000$, $v_P = v_T = v_R = 2 * 10^{-9} (\mu m)^3$, $f_R = 0.4, K_Z = 10^{-8} hr^{-1} (\mu m)^3$, $Z_c = 60$, $Z_r = 60$, $\tau_1 = 0$. The stochastic trajectory (not shown) for parameter set 2 is qualitatively similar to that in Fig. 2A but with completely different averages in the statistical steady state: $\langle P \rangle = 2.766 * 10^4, \langle T \rangle = 4.489 * 10^4, \langle R \rangle = 1.197 * 10^3, \langle V_b \rangle = 1.475 * 10^{-4} (\mu m)^3, \langle V_d \rangle = 2.953 * 10^{-4} (\mu m)^3, \langle \Delta \rangle = 1.478 * 10^{-4} (\mu m)^3, \langle \tau \rangle = 9.258 * 10^{-3} hr, \langle \alpha \rangle = 74.975 hr^{-1}$. Also given are the same distributions for two other models, the XZ model and the XYZ model defined in section IV E. XZ model parameters: $\mu = 10, K_Z = 10^{-3}, v = 1$. XYZ model parameters: $c_1 = 10, c_2 = 1, K_Z = 10^{-3}, v_1 = v_2 = 1$. For both models $Z_c = 40, Z_r = 20, \tau_1 = 0$. It is seen that the rescaled $\Delta$, $\tau$ and $V_d$ distributions are robust to parameter changes in the PTRZ model and even to changes in the model itself (as long as $N = Z_c - Z_r/2$ is the same), whereas the rescaled $\alpha$ distribution is not. All the models are run with intrinsic stochasticity in the dynamics of all the chemicals and no other source of stochasticity.

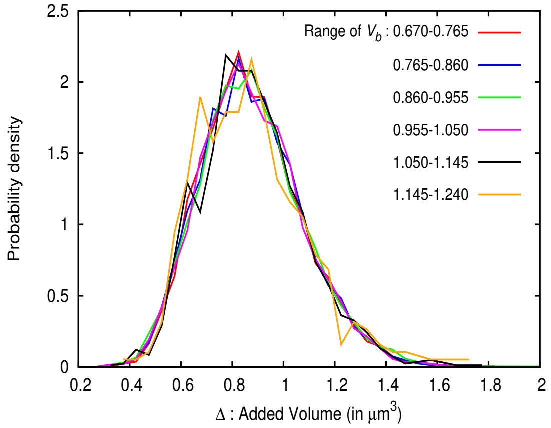


FIG. S2: **Collapse of the conditional $\Delta$ distributions for different $V_b$ in the PTRZ model**. The probability distributions of $\Delta$ for different (binned) values of $V_b$ are shown. Their collapse onto each other shows that the distribution of the volume added in a generation is independent of the birth volume, a strong test of the adder property. Parameter values are the same as in Fig. 1.

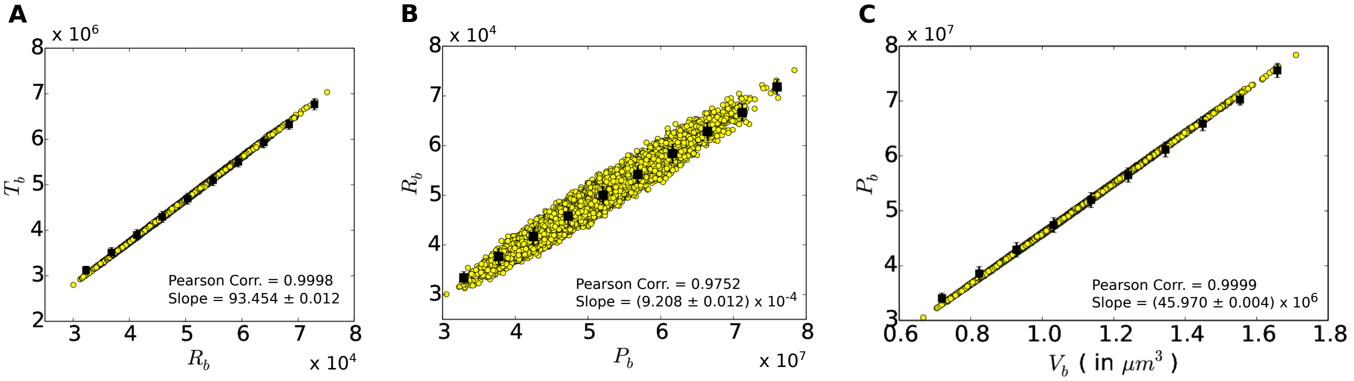


FIG. S3: **Extensive quantities are linearly correlated with each other in the PTRZ model**. Scatter plots of intracellular populations and volume of the PTRZ cell at birth for $10^4$ cells. The correlations are close to unity. The slopes of the best fit lines of the binned data are close to the ratios expected from Eqs. (8a), (8b), (3), namely 93.5, $9.64 \times 10^{-4}$, and $4.58 \times 10^7$.

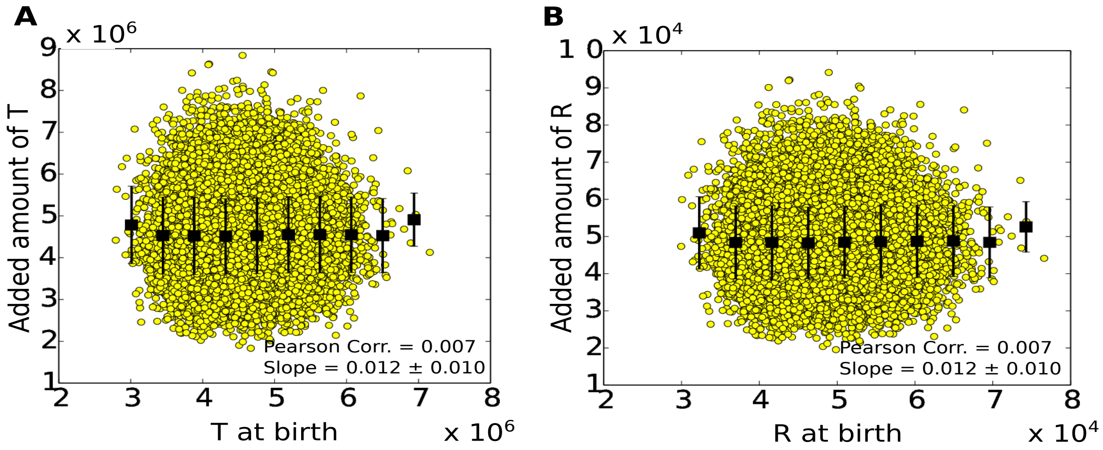


FIG. S4: **The adder property for intracellular populations in the PTRZ model**. **A**(**B**) The scatter plots for added population of $T$ ($R$) in a generation versus $T$ ($R$) at the beginning of the generation. The figures show that the increments in $T$ and $R$ from birth to division are essentially uncorrelated with the values at birth.

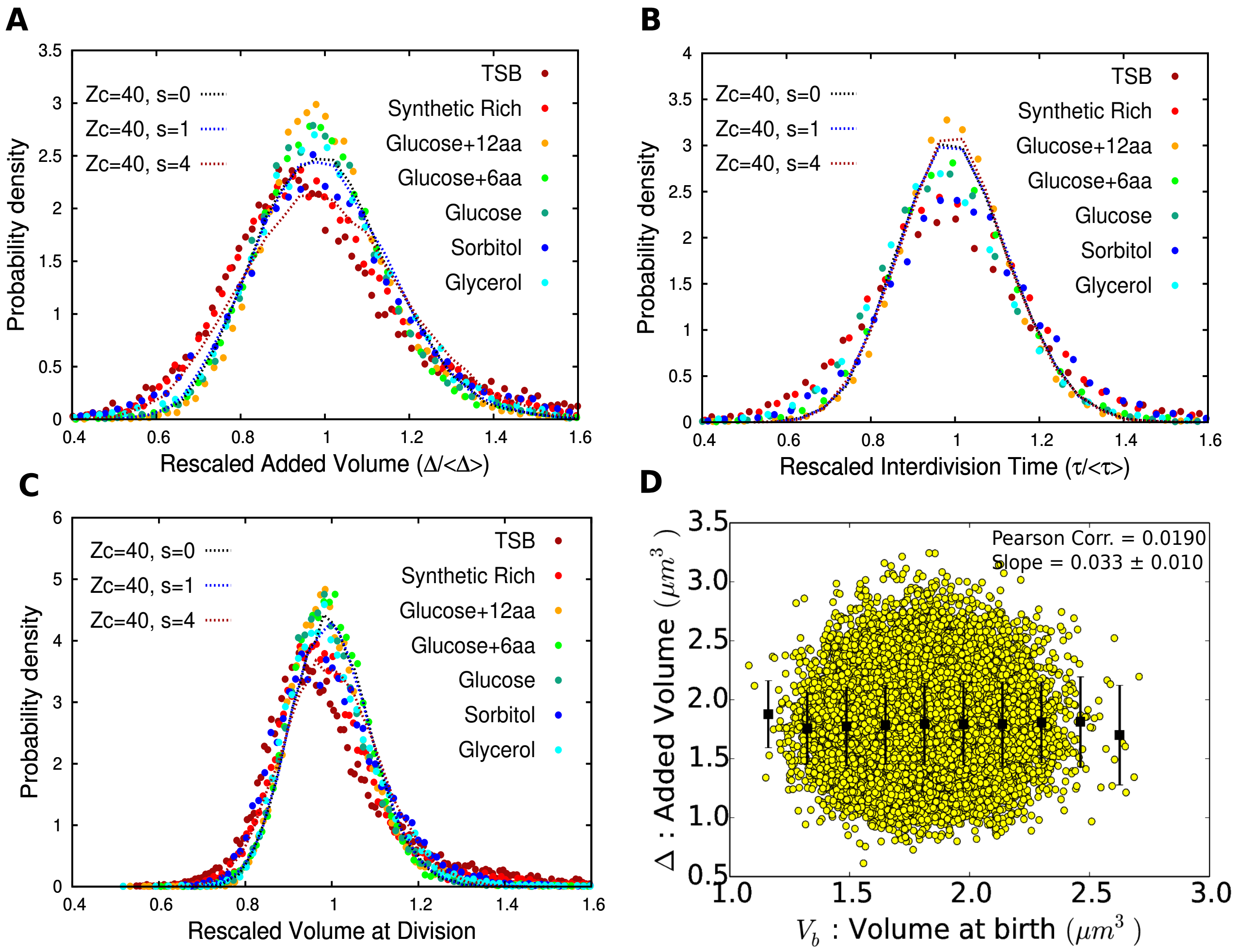


FIG. S5: **Effect of including partitioning stochasticity in PTRZ model.** Simulations are done as described in the legend of Fig. 6. $s = 0$ curves mean that the only stochasticity is the intrinsic stochasticity in the chemical dynamics of $P, T, R, Z$ and $s > 0$ means that partitioning stochasticity is also included as described in section III B 5. **A,B,C** display the distributions of rescaled $\Delta, \tau, V_d$. It is seen that at $s = 1$ partitioning stochasticity does not significantly alter these three distributions. At $s = 4$ the $\tau$ distribution is still unaltered while the distributions of $\Delta$ and $V_d$ broaden slightly. This is in sharp contrast to the $\alpha$ distribution which broadens quite significantly at $s = 4$ (Fig. 6D). Data points from the experiment of [11] are shown for reference and are the same as in Fig. 6. **D** Scatter plot of $\Delta$ vs $V_b$ for $Z_c = 40$, $s = 4$, showing that the adder property of the volume is preserved at this strength of partitioning stochasticity.

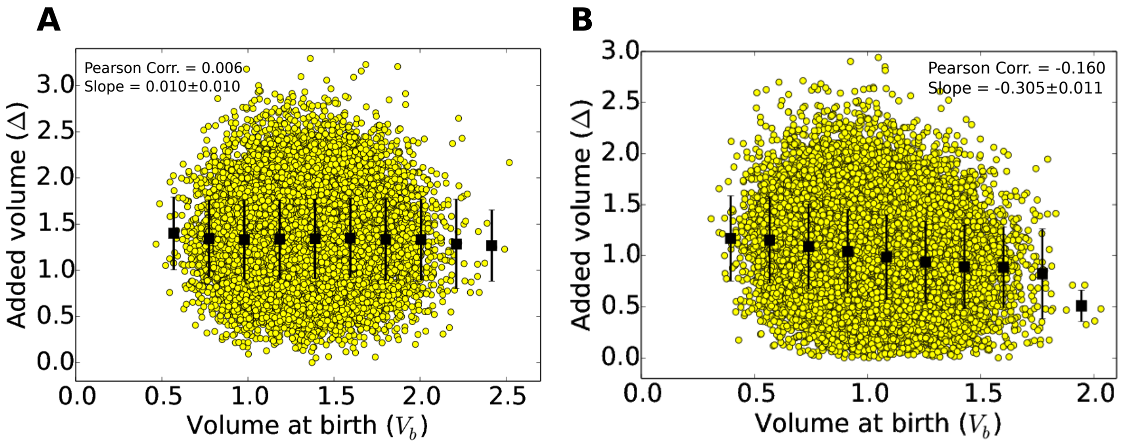


FIG. S6: **Effect of replacing a sharp cutoff $Z_c$ by a stochastic cutoff**. Simulations are done with stochasticity in the chemical dynamics of PTRZ and stochasticity in the value of the division threshold as discussed in section III B 5. At each generation the value of the threshold $Z'_c$ at which division is triggered is independently chosen from a gaussian distribution with a mean $= Z_c = 30$ and s.d. $= s' = 8$. There is no partitioning stochasticity. **A** $Z_r = 0$. The adder property of the cell volume is preserved as evidenced by the slope of mean $\Delta$ vs $V_b$ being essentially zero. **B** $Z_r = Z'_c/2$. The adder property of the cell volume is lost as the slope of mean $\Delta$ departs significantly from zero.

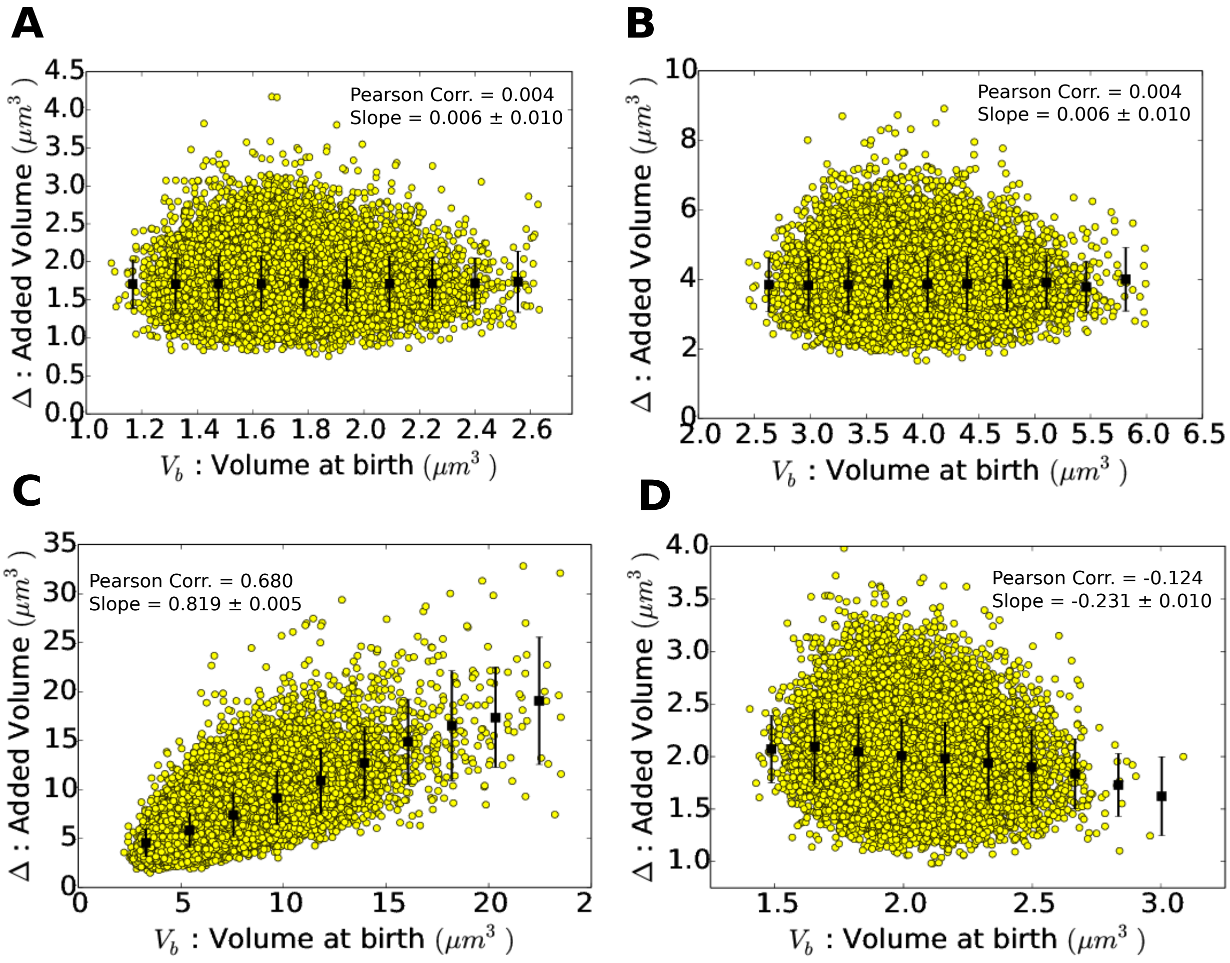


FIG. S7: **Effect of auto-regulation of $Z$ on the adder property of cell volume**. **A, B** When the regulatory function $r$ in Eq. (21) is a function of $Z$ alone, the adder property is preserved (**A** positive auto-regulation; **B** negative auto-regulation). **C,D** When the regulatory function $r$ is a function of $Z/V$, the adder property is lost (**C** positive auto-regulation; **D** negative auto-regulation). Simulations are done with stochasticity in the chemical dynamics of PTR and Z and no other source of stochasticity. Model parameters are as in Fig. 1, except $Z_c = 25$, and $Z_r = 10$. The regulatory function is given by **A**: $r(Z) = Z/(K+Z)$; **B**: $r(Z) = K/(K+Z)$; **C**: $r(P,T,R,Z) = (Z/V)/(K+(Z/V))$; **D**: $r(P,T,R,Z) = K/(K+(Z/V))$, where $V$ is given by Eq. (3) of the main paper. $K = 7$ in **A** and **B**, and $K = 7\ (\mu m)^{-3}$ in **C** and **D**.

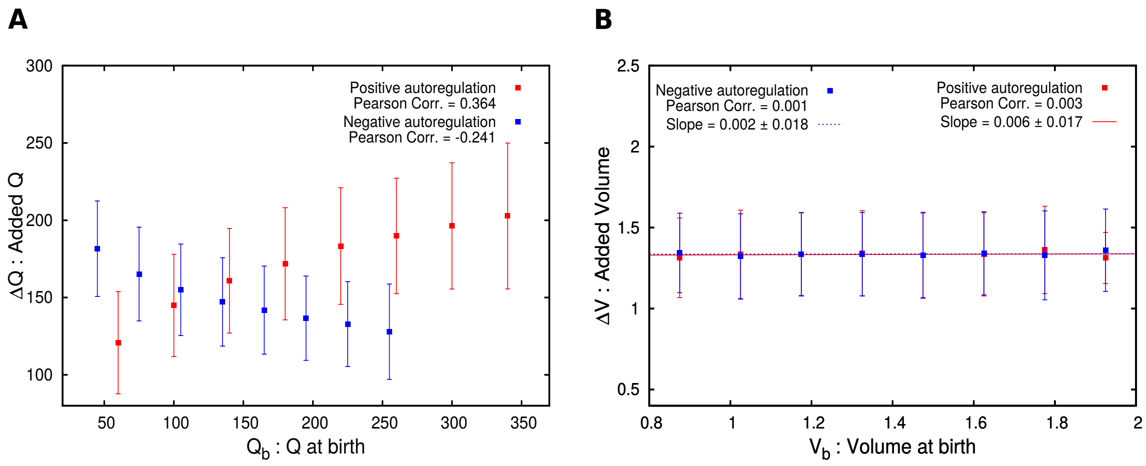


FIG. S8: **Example of a molecular population $Q$ that does not show the adder property while $V$ does.** **A** The increment in $Q$ in a generation, $\Delta Q$, is positively correlated with the birth value $Q_b$ when $Q$ is positively auto-regulated, and negatively when $Q$ is an auto-repressor. **B** $V$ satisfies the adder property in both cases. Simulations are done with stochasticity in the chemical dynamics of PTRZ and Q as well as partitioning stochasticity in the populations. The partitioning stochasticity parameter $s$ equals 2 for the PTR populations and 3 for the Q population. Model parameters are as in Fig. 1, with $K_Q = 10^{-10} hr^{-1} (\mu m)^3$ in eqn. (22), $K = 100\ (\mu m)^{-3}$, $h' = 1$.

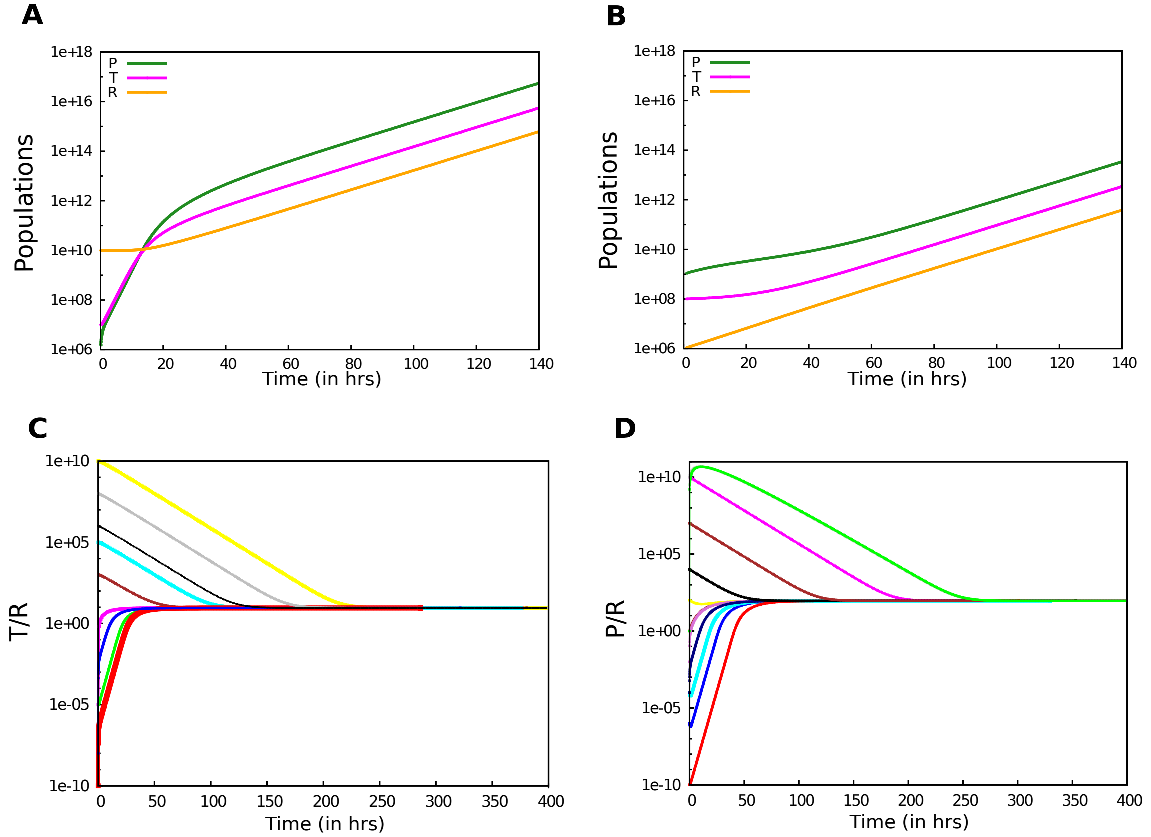


FIG. S9: **The PTR model without division has an exponentially growing attractor**. Simulations are shown for a system in which there are only three chemical populations $P, T, R$ satisfying Eqs. (2) with $V$ given by (3). The population $Z$ is absent and there is no division control. The populations increase indefinitely with time. Eqs. (2) were simulated with the following numerical values of the parameters: $K_P = 1$, $k = 1$, $m_T = m_R = 1$, $d_T = d_R = 0$, $f_R = 0.1$, $v_P = v_T = v_R = 1$. **A** and **B** plot $P$, $T$ and $R$ vs $t$ in a semi-log plot, starting from two sets of initial conditions, **A** IC1: $P_0 = 10^6$, $T_0 = 10^7$, $R_0 = 10^{10}$ and **B** IC2: $P_0 = 10^9$, $T_0 = 10^8$, $R_0 = 10^6$. The straight lines with the same slope for all the three chemical species imply an asymptotic exponential growth consistent with (7). The value of $\mu = 0.09$ obtained numerically from the slope agrees with the formula (8c) for these parameter values. **C** $T/R$ and **D** $P/R$ as functions of time starting from a wide range of initial conditions. $1311 = 11^3$ simulations were done in which each of $P(0)$, $T(0)$ and $R(0)$ took values from the set $\{10^0, 10^1, 10^2, \ldots\ldots, 10^{10}\}$. A subset of the simulations is shown. The ratios always converge to the values $T/R = 9.0$ and $P/R = 90.0$, which are the same as those predicted by (8a) and (8b). This shows that an exponentially growing trajectory (7) along the curve of balanced growth defined by (8) is a stable attractor of the dynamics (and possibly a global attractor) for these parameter values indicating that this is a generic property of the PTR model. This was repeated for other parameter values and the same behaviour was observed whenever the r.h.s. of (8c) was positive. (Specifically the four parameters $K_P, k, m_T, m_R$ were each independently changed by a factor of $10^{-2}$ or $10^2$ from their values mentioned above keeping the other parameters fixed, giving a total of $2^4 = 16$ parameter sets.)

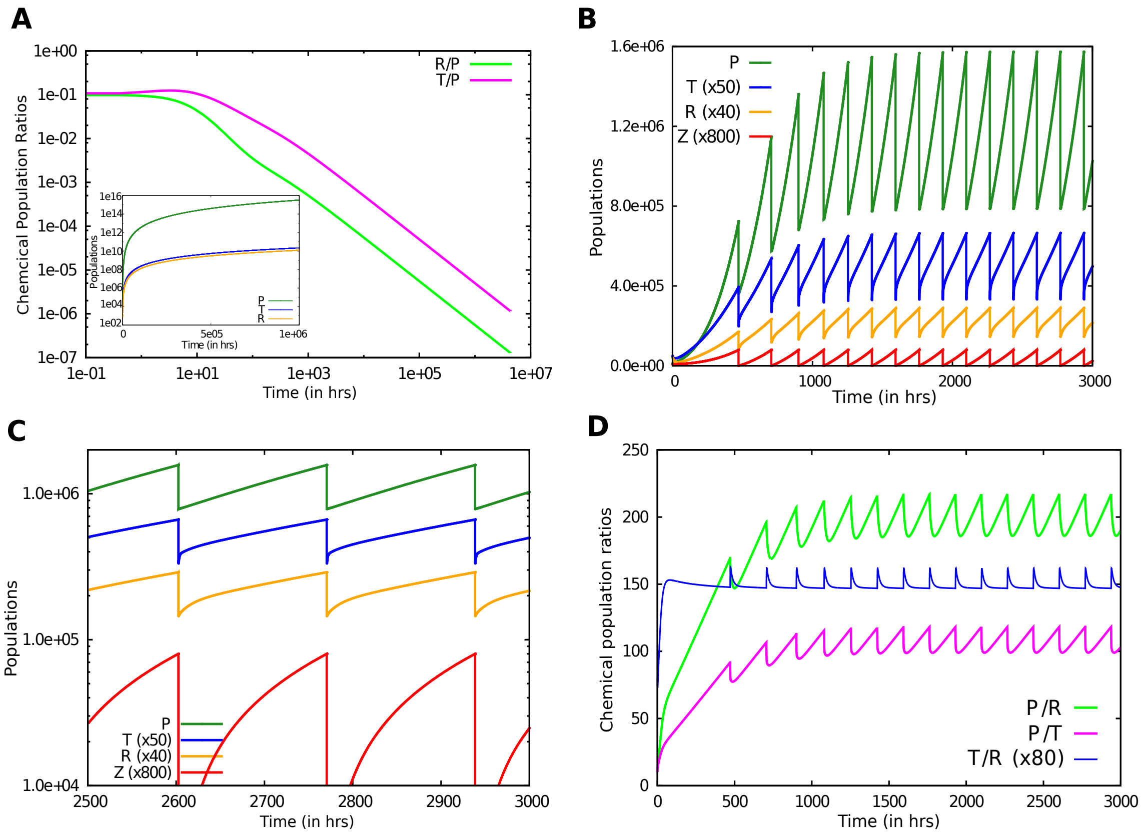


FIG. S10: **A non-class-I system shows departures from exponential growth**. Simulations are shown for a system with the same chemical rate equations as for the PTR cell (Eq. (2)), but with $V = wS(t)^{\gamma}$ and $S = aT$, where $w, a, \gamma$ are constants. **A** Ratios $T/P$ and $R/P$ vs time for an indefinitely growing PTR-II cell in log-log axes, with absolute populations in the inset. The ratios do not become constant as in S9C,D but instead asymptotically decline as a negative power of $t$ showing that (7) is not an attractor for the system. Eventually the $P$ population completely overshadows $T$ and $R$. Parameters: $K_P = k = m_T = m_R = w = a = 1$, $f_R = 0.1$, $d_T = 0.5$, $d_R = 0.1$, $\gamma = 3/2$. **B** A trajectory of the same system but with division implemented through the $Z$ sector as for the PTRZ model ($K_Z = 10^{-4}, Z_c = 100, Z_r = 0, \tau_1 = 0$). Division produces a nontrivial limit cycle attractor in which all three chemicals survive. **C** Log of the populations vs time in the limit cycle attractor. Note the departure from linearity in the growth phase. **D** Ratios $T/P$ and $R/P$ vs time for a trajectory of the system. The ratios continue to oscillate after reaching the attractor, showing that between birth and division the trajectory does not satisfy the exponential ansatz Eq. (7) (compare with the second inset of Fig. 1B where they become constant in the attractor). The initial populations at $t = 0$ are $P = 10^4, T = 10^3, R = 10^3, Z = 9$ in the simulations shown above. The long time behaviour in this figure is reproduced for diverse initial conditions, and the above mentioned qualitative character is numerically observed to be robust to parameter variations of the system (details not shown).

### S2. SUPPLEMENTARY TEXT

#### A. Cell volume at birth in the limit cycle attractor of the PTRZ model when $\tau > \tau_1 > 0$

Let us denote by $t = 0$ the time of birth of the daughter cell, and by $t = \tau$ the time of division. On the limit cycle attractor $\tau$ is fixed by the parameters of the PTR sector and given by (9). Since all populations including $Z$ are halved at division, $Z_b \equiv Z(0) = (1/2)Z(\tau) \equiv (1/2)Z_d$. By assumption, $Z_b < Z_c$. $Z(t)$ increases with time following the differential equation (10) until at a time denoted by $t_1$ it reaches $Z_c$. Thus in the period between 0 and $t_1$, $Z(t)$ is given by (12). Thus $Z_c = Z_b + B(e^{\mu t_1} - 1)$ or

$$Z_b = Z_c - B(e^{\mu t_1} - 1). \tag{S1}$$

$Z$ is instantaneously reset to $Z_r$ at $t_1$. In the time between $t_1$ and $\tau$ (and this interval is precisely $\tau_1$ by assumption: $\tau_1 = \tau - t_1$), $Z$ increases from the value $Z_r$ to $Z_d$ again following the equation (10). Note that $\dot{Z}$ is continuous in the interval between birth and division because the $X$ sector populations evolve continuously in this period, even though $Z$ has a discontinuity at $t_1$. The solution for $Z(t)$ in the period between $t_1$ and $\tau$ is $Z(t) = Z_r + B(e^{\mu t} - e^{\mu t_1})$. Hence $Z(\tau) = Z_r + B(e^{\mu\tau} - e^{\mu t_1})$ or

$$Z_b = (1/2)Z(\tau) = (1/2)[Z_r - B(e^{\mu\tau} - e^{\mu t_1})]. \tag{S2}$$

Equating (S1) and (S2) to eliminate $Z_b$, and using $t_1 = \tau - \tau_1$, $e^{\mu\tau} = 2$, we get

$$B = (Z_c - \frac{Z_r}{2})e^{\mu\tau_1}. \tag{S3}$$

The formula (14) for $V_b$ then follows immediately from the relation $C = \mu B$ and (11).

#### B. Distribution of the first passage time

For completeness we give a derivation of the first passage time distribution. Consider first the deterministic dynamics

$$\dot{Z} = H(t), \tag{S4}$$

where $H(t)$, the rate of the reaction in which $Z$ is produced, is a given non-negative function of time. This has the solution

$$Z(t) = Z_b + \lambda(t), \quad \lambda(t) = \int_0^t dt' H(t'), \tag{S5}$$

where $Z_b$ is the value of $Z$ at $t = 0$. The interdivision time $\tau$ is the solution of $\lambda(\tau) = Z_c - Z_b \equiv N$. For the specific case $H(t) = Ce^{\mu t}$ appearing in (10), the deterministic solution is given in (12) and

$$\tau = \frac{1}{\mu} \ln{(1 + \frac{\mu N}{C})}. \tag{S6}$$

Now consider the case where the dynamics of $Z$ is the stochastic version of (S4). That is, $H(t)$ is the instantaneous probability of occurrence, per unit time, of the reaction in which a single molecule of $Z$ is produced. Thus the probability that $Z$ increases by unity in the small time interval $(t, t + \delta t)$ is given by $H(t)\delta t$, and the probability that it remains unchanged is $1 - H(t)\delta t$. Let the probability at time $t$ that the population of this molecular species is the integer $Z$ be given by $P(Z, t)$. Then $P(Z, t + \delta t) = P(Z - 1, t)H(t)\delta t + P(Z, t)(1 - H(t)\delta t)$. It follows that $P$ satisfies the master equation

$$\frac{\partial}{\partial t} P(Z, t) = H(t)[P(Z - 1, t) - P(Z, t)]. \tag{S7}$$

The relevant solution to this for $t \geq 0$ is given by

$$P(Z, t) = \begin{cases} 0 & \text{if } Z = 0, 1, \ldots, Z_b - 1, \\ \lambda^{Z - Z_b} e^{-\lambda}/(Z - Z_b)! & \text{if } Z = Z_b, Z_b + 1, \ldots, \end{cases} \tag{S8}$$

where $\lambda = \lambda(t)$ is given in (S5). This solution satisfies the required initial condition $P(Z, 0) = \delta_{Z,Z_b}$. It is the Poisson distribution (shifted from the origin by an amount $Z_b$), with a mean $\langle Z \rangle(t)$ given by $Z_b + \lambda(t)$, the same as the deterministic solution (S5). For any particular stochastic trajectory in the ensemble (we are considering the ensemble of trajectories that start at $t = 0$ with a fixed $Z = Z_b$ and have a fixed probability density function $H(t)$ for $t \geq 0$), the time to division, $\tau$, is the first passage time for $Z$ to reach $Z_c$. Since $Z$ is a non-decreasing quantity, the first passage time distribution is given by $\mathcal{P}(\tau) = \frac{d}{d\tau} S(Z_c, \tau)$, where $S(Z_c, \tau) = \sum_{Z=Z_c}^{\infty} P(Z, \tau)$. (This follows from the fact that the fraction of trajectories that have reached or passed $Z_c$ by a time $\tau$ is the probability that $Z \geq Z_c$ at time $\tau$, namely, $\sum_{Z=Z_c}^{\infty} P(Z, \tau)$. Thus the fraction of trajectories reaching $Z_c$ in the time interval $(\tau, \tau + d\tau)$, $\mathcal{P}(\tau)d\tau$, is given by $S(Z_c, \tau + d\tau) - S(Z_c, \tau)$). Differentiating (S8) with respect to $t$ and taking the sum over $Z$ yields the result

$$\mathcal{P}(\tau) = \frac{\lambda^{N-1} e^{-\lambda}}{(N - 1)!} \frac{d\lambda}{d\tau}, \tag{S9}$$

where $\lambda \equiv \lambda(\tau) = \int_0^\tau dt' H(t')$. This proves Eq. (15) for a more general setting than that mentioned in the main text (in which $Ce^{\mu t}$ is replaced by a general non-negative function $H(t)$).

#### C. Effect of $Z$ autoregulation on the first passage time distribution and the adder property of cell volume

In this section we show that the adder property of cell volume in the PTRZ model is preserved when the $Z$ dynamics is autoregulated via a nonlinear function $r(Z)$. In other words we assume that the dynamics of $Z$ is the stochastic version of Eq. (21),

$$\dot{Z} = H(t)r(Z), \tag{S10}$$

where $r(Z)$ depends nontrivially on $Z$. Further,

$$H(t) = Ce^{\mu t}, \quad C = c_1 V_b, \quad V(t) = V_b e^{\mu t}, \tag{S11}$$

where $c_1$ is a constant independent of generation and hence of $V_b$ (see, e.g., Eq. (11)). This encapulates the approximation that the dynamics of $X$ sector chemicals (e.g., $P$, $T$, and $R$) can be treated deterministically and the $X$ sector has settled down into its attractor on the curve of balanced growth. Therefore in any generation, $C$ is proportional to the birth volume $V_b$ in that generation with a proportionality constant that is the same for all generations. Further the cell volume grows exponentially with time (see remarks at the beginning of Section IV D of the main text).

Consider first the deterministic evolution of $Z$ under Eq. (S10). The equation can be integrated to give

$$g(Z) = \frac{C}{\mu}(e^{\mu t} - 1), \quad g(Z) \equiv \int_{Z_b}^{Z} \frac{dZ'}{r(Z')}, \tag{S12}$$

where $Z_b$ is the value of $Z$ at birth ($t = 0$). Thus the division volume is given by

$$V_d = V_b e^{\mu\tau}, \tag{S13}$$

where $\tau$, the time of division, is given by the solution of

$$g(Z_c) = \frac{c_1 V_b}{\mu}(e^{\mu\tau} - 1). \tag{S14}$$

Thus the added volume is given by

$$\Delta \equiv V_d - V_b = V_b(e^{\mu\tau} - 1) = \frac{\mu g(Z_c)}{c_1}. \tag{S15}$$

If $r(Z)$ is dependent on $Z$ alone (and not on any other dynamical variable $X_i$ or the volume $V$), the quantity $g(Z_c)$ obtained from the integral in (S12) is independent of $V_b$. Then $\Delta$ is independent of $V_b$ because $c_1$ is independent of $V_b$. Note that $\tau$ given by (S14) and $V_d$ given by (S13) are not independent of $V_b$, but $\Delta$ is. However when $r$ depends on other variables in addition to $Z$, the above independence of $\Delta$ with respect to $V_b$ fails in general. E.g., if $r$ is a function of $Z/V$ instead of $Z$ (see examples in the caption of Figs. S7C and S7D), then $g(Z_c)$ and hence $\Delta$ start to depend upon $V_b$.

We now consider the stochastic evolution of $Z$ under the assumption that the probability of producing a $Z$ molecule at any time is proportional to the r.h.s. of (S10) and show that the resulting distribution of $\Delta$ is independent of $V_b$ when $r$ depends on $Z$ alone. The proof proceeds by showing that in every increment of $Z$ by unity the distribution of added volume is independent of the volume at the previous value of $Z$. Therefore the distribution of the total added volume in the $N$ unit increments of $Z$ starting from $Z_b$ is independent of the starting volume $V_b$. Consider the ensemble of trajectories that start with $Z = Z_b$ and $V = V_b$ at $t = 0$. In this ensemble $V(t) = V_b e^{\mu t}$ but $Z$ is stochastic. For any given stochastic trajectory of the ensemble, define $t_i$ as the time when $Z$ jumps from $Z_b + (i-1)$ to $Z_b + i$, $i = 1, \ldots, N$ with $t_0 \equiv 0$. Further, $u_i \equiv t_i - t_{i-1}$, $V_i \equiv V(t_i)$, and $v_i \equiv V(t_i) - V(t_{i-1})$, $i = 1, \ldots, N$. $u_i$ is by definition the time taken for $Z$ to jump from $Z + (i-1)$ to $Z + i$. $u_i$ are the basic stochastic variables of the process, in terms of which $t_i$, $V_i$, $v_i$ and $\Delta$ are determined. $t_i = \sum_{j=1}^{i} u_j$, $V_i = V_b e^{\mu t_i}$, $v_i = V_b(e^{\mu t_i} - e^{\mu t_{i-1}})$, $\Delta = \sum_{i=1}^{N} v_i$.

Say at some time $t' > 0$, $Z$ equals $Z'$. After a certain time interval, which is stochastic, $Z$ will become $Z' + 1$. Consider the probability distribution of this time interval, denoted $u$. Specifically, let $\mathcal{P}(u|Z', t')du$ be the probability that $Z$ jumps from $Z'$ to $Z' + 1$ in the time interval $t' + u$ to $t' + u + du$, given that at time $t'$, $Z$ was equal to $Z'$. Then we find that

$$\mathcal{P}(u|Z', t') = e^{-\lambda(Z', V_b, t'+u, t')} \frac{\partial}{\partial u} \lambda(Z', V_b, t' + u, t') \tag{S16}$$

where

$$\lambda(Z, V_b, t, t') \equiv \int_{t'}^{t} H(t'')r(Z)dt'' = a(Z)V_b(e^{\mu t} - e^{\mu t'}),$$
$$a(Z) \equiv \frac{c_1 r(Z)}{\mu}, \tag{S17}$$

and in taking the partial derivative with respect to $u$, $Z'$, $V_b$ and $t'$ are held constant. The proof of (S16) is given later. For now, we use (S16) to derive the distribution of the added volume. Let $v$ denote the volume increment in the same time interval $t'$ to $t' + u$. Then

$$v = V(t' + u) - V(t') = V_b(e^{\mu(t'+u)} - e^{\mu t'}) = \frac{\lambda(Z', V_b, t' + u, t')}{a(Z')} \tag{S18}$$

Given $V_b$, $Z'$ and $t'$, $v$ is completely determined by $u$. Hence the probability distribution of $v$ (denoted $\mathcal{R}(v)$) is determined by the probability distribution of $u$. More precisely, let $\mathcal{R}(v|Z', t')dv$ be the probability that the volume increment is in the range $v$ to $v + dv$ between the time $t'$ and the time when $Z$ jumps from $Z'$ to $Z' + 1$, given that $Z$ equalled $Z'$ at time $t'$. By definition $\mathcal{R}(v|Z', t')dv = \mathcal{P}(u|Z', t')du$. Therefore $\mathcal{R}(v|Z', t') = \mathcal{P}(u|Z', t')\frac{\partial u}{\partial v}$, where, in the partial derivative $\frac{\partial u}{\partial v}$ all quantities other than $v$ on which $u$ depends (such as $Z'$, $t'$, $V_b$) are held fixed. Therefore

$$\mathcal{R}(v|Z', t') = e^{-\lambda}\frac{\partial \lambda}{\partial u}\frac{\partial u}{\partial v} = e^{-\lambda}\frac{\partial \lambda}{\partial v} = e^{-a(Z')v}a(Z'), \tag{S19}$$

where we have used (S16) in the first step and (S18) in the last.

The key point is that the probability distribution of the volume increment $v$, $\mathcal{R}(v|Z', t')$ depends only on $v$ and $Z'$

and not on $V_b$ or $t'$, while $\mathcal{P}$ the probability distribution of $u$ (the time taken for $Z$ to increment by unity) depends not only on $u$ and $Z'$ but also on $V_b$ and $t'$ (see Eq. (S16)). Thus the probability distribution of $u_1$ in the ensemble of trajectories characterized by $Z_b$ and $V_b$ is

$$\begin{aligned}\mathcal{P}_1(u_1|Z_b,0') &= e^{-\lambda(Z_b,V_b,u,0)}\frac{\partial}{\partial u}\lambda(Z_b,V_b,u,0)\\ &= e^{-\lambda(Z_b,V_b,u,0)}c_1 r(Z_b)V_b,\end{aligned} \tag{S20}$$

while the distribution of $v_1$ is

$$\mathcal{R}_1(v_1|Z_b,0) = e^{-a(Z_b)v_1}a(Z_b). \tag{S21}$$

The probability distribution of $u_2$ is conditional on the value of $u_1$: One can consider the restricted ensemble of trajectories that start at $t=0$ with $Z=Z_b$ and $V=V_b$ AND for which $Z$ jumps to $Z_b+1$ at $t=u_1$. In this ensemble the probability of $Z$ jumping from $Z_b+1$ to $Z_b+2$ in the time interval $u_1+u_2$ to $u_1+u_2+du_2$ is

$$\begin{aligned}&\mathcal{P}_2(u_2|Z_b,V_b,u_1)du_2 = \mathcal{P}(u_2|Z_b+1,u_1)du_2\\ &= e^{-\lambda(Z_b+1,V_b,u_1+u_2,u_1)}\frac{\partial}{\partial u_2}\lambda(Z_b+1,V_b,u_1+u_2,u_1).\end{aligned} \tag{S22}$$

The probability that in the same ensemble the added volume in the interval in which $Z$ jumps from $Z_b+1$ to $Z_b+2$ lies between $v_2$ and $v_2+dv_2$, defined as $\mathcal{R}_2(v_2|Z_b,V_b,u_1)dv_2$, equals $\mathcal{R}(v_2|Z_b,u_1)dv_2$. Thus

$$\mathcal{R}_2(v_2|Z_b,V_b,u_1) = e^{a(Z_b+1)v_2}a(Z_b+1), \tag{S23}$$

which is independent of $V_b$ and $u_1$, and hence of $v_1$.

Hence the joint probability of the first volume increment lying between $v_1$ and $v_1+dv_1$ and second increment lying between $v_2$ and $v_2+dv_2$ is the product,

$$\begin{aligned}&\mathcal{R}^{(2)}(v_1,v_2|Z_b,V_b)dv_1dv_2\\ &\quad= e^{-a(Z_b)v_1}a(Z_b)dv_1\; e^{-a(Z_b+1)v_2}a(Z_b+1)dv_2.\end{aligned} \tag{S24}$$

The same arguments can be repeated with the third increment and so on; all successive volume increments are independent of the previous ones. Therefore the joint probability of $N$ increments is

$$\begin{aligned}&\mathcal{R}^{(N)}(v_1,\dots,v_N|Z_b,V_b)dv_1dv_2dv_3...dv_N\\ &\quad= \prod_{i=1}^{N}(e^{-a(Z_b+i-1)v_i}a(Z_b+i-1)dv_i).\end{aligned} \tag{S25}$$

The above distribution is also independent of $V_b$ because all the component distributions are.

The distribution of the total added volume is then given by

$$\mathcal{R}(\Delta)d\Delta = \int_{\text{Shell}(\Delta,d\Delta)}\mathcal{R}^{(N)}(v_1,\dots,v_N|Z_b,V_b)dv_1...dv_N, \tag{S26}$$

where Shell$(\Delta,d\Delta)$ is the $N$-dimensional region in the positive orthant of the $v_1,v_2,\dots,v_N$ space between the hypersurfaces $v_1+\ldots+v_N=\Delta$ and $v_1+\ldots+v_N=\Delta+d\Delta$. Since $\mathcal{R}^{(N)}$ is independent of $V_b$, so is $\mathcal{R}(\Delta)$; this proves the adder property. As a special case when $r(Z)=1$, the integrand in (S26) depends only on $\Delta$, hence the integral equals $e^{-a\Delta}\times$(Volume of Shell$(\Delta,\mathrm{d}\Delta)$), and (S26) reduces to the old distribution for the added volume distribution given by (16).

Proof of (S16): By definition, $\mathcal{P}(u|Z',t')du=$ [Probability that $Z$ remains equal to $Z'$ up to time $t'+u$ given that $Z$ equalled $Z'$ at the earlier time $t'$] $\times$ [probability that $Z$ jumps from $Z'$ to $Z'+1$ in the time interval $t'+u$ to $t'+u+du$]. Thus

$$\mathcal{P}(u|Z',t')du = P(Z',t'+u|Z',t')H(t+u)r(Z')du, \tag{S27}$$

where $P(Z',t|Z',t')\equiv$ Probability that $Z$ remains equal to $Z'$ at time $t$ given that $Z$ equalled $Z'$ at an earlier time $t'$. ($H(t)r(Z')dt$ is the probability that $Z$ jumps from $Z'$ to $Z'+1$ in the time interval $t$ to $t+dt$.) From the definition of $P$ it follows that

$$P(Z',t+dt|Z',t') = P(Z',t|Z',t')[1-H(t)r(Z')dt], \tag{S28}$$

$[1-H(t)r(Z')dt]$ being the probability that $Z$ does not jump from $Z'$ to $Z'+1$ in the time interval $t$ to $t+dt$. This implies that $P$ satisfies the differential equation $\frac{\partial}{\partial t}P(Z,t|Z,t')=-H(t)r(Z)$ whose solution is

$$P(Z,t|Z,t') = e^{-\lambda(Z,V_b,t,t')}, \tag{S29}$$

where $\lambda$ is given by (S17). Substituting (S29) in (S27) and using (S17) gives (S16).

The above proofs are valid for a much larger class of models than the PTRZ model when the functions $f_i(\mathbf{X})$ and $h(\mathbf{X},Z)$ appearing in Eq. (1) are homogeneous degree one functions of the $X_i$, since (S10) and (S11) hold for the general case also, as discussed in Section V C 1 of the main text.

#### D. Remarks about solutions of (27)

It was shown in the main text that for class-I systems (that satisfy (24)), the exponential ansatz (25) is a solution of the dynamics (1a) if and only if (27) or equivalently (28) holds. Equation (28) is a set of $n$ equations for the $n$ unknowns $\psi_i$ ($i=1,2,\dots,n-1$) and $\mu$. Generically such a system has a finite set of solutions because the number of unknowns equals the number of equations. (For non-generic systems whose parameters have been fine-tuned we can get a one (or more) parameter family of solutions.) In the PTR model, two solutions were found [50], of which one turned out to be non-physical (it gave negative populations). That left a unique solution of (27) which is given in (8). In chemical dynamics the functions $f_i$ that appear are typically algebraic functions. Thus the ratios of chemicals and the growth rate for exponential solutions of class-I systems are determined as solutions of a system of $n$ coupled algebraic equations.

Consider the $n$-dimensional phase space $\Gamma$ whose coordinates are $X_i$. A generic point $\mathbf{X} = (X_1, X_2, \dots X_n)$ in $\Gamma$ specifies all the chemical populations. Suppose the system is class-I and we have a solution $(\mu, \mathbf{X})$ to (27). We consider only non-trivial solutions (in which $\mathbf{X}$ is not identically $\mathbf{0}$). Then without loss of generality we can consider $X_n \neq 0$. Defining $\psi_i = X_i/X_n$, we have that $(\mu, \boldsymbol{\psi})$ is a solution to (28), where $\boldsymbol{\psi} = (\psi_1, \dots, \psi_n) \equiv (X_1/X_n, \dots, X_{n-1}/X_n, 1)$. The vector $\boldsymbol{\psi}$ specifies the direction of a straight line passing through the origin in the $n$-dimensional phase space $\Gamma$. Consider a trajectory starting at time $t = 0$ on this line, i.e., $\mathbf{X}(0) = c\boldsymbol{\psi}$ for some fixed constant $c$. Then since $(\mu, \mathbf{X}(0))$ satisfies (27), (25) is a solution to (1a). In this exponentially growing solution the ratios $X_i(t)/X_n(t)$ are time independent. Therefore the trajectory stays on the same straight line as time progresses. In other words the line is an invariant manifold of the dynamics. In the main text this line is referred to as a 'curve of balanced growth' (CBG) for the system, because ratios of populations remain constant if the system starts on this curve. If the system is not class-I, in general one cannot get exponentially growing solutions to equation (1a).

To be physically acceptable as ratios of populations, the components $\psi_1$, $\psi_2$, $\dots \psi_{n-1}$ of the solution of (28) must be real and non-negative. For a growing cell the growth rate $\mu = f_n(\psi_1, \psi_2, \dots, \psi_{n-1})$ should also be real and positive. In addition the exponential solution should also be a stable attractor. In the PTR case the analytic formulae (8) themselves provide the information about the parameter domain where the physical quantities have the right sign. Our numerical investigation suggests (see caption of Fig. S9) that there is a substantial parameter domain where it is also an attractor. However in general the class-I condition by itself does not guarantee that a physically acceptable exponentially growing solution does exist or that it is a stable attractor. One can construct examples of class-I systems for which a physically acceptable exponentially growing solution does not exist, or exists but is unstable. Nevertheless for all the class-I systems that we have studied motivated by cellular dynamics (which has an underlying autocatalytic network of chemical reactions) we have found physically valid exponential solutions that are stable attractors in a substantial parameter domain. These examples will be discussed elsewhere.
